# Old, broad-leaved stands support both high biodiversity and carbon storage in German forests

**DOI:** 10.1101/2024.02.06.578731

**Authors:** K. Springer, P. Manning, A.L. Boesing, C. Ammer, A.M. Fiore-Donno, M. Fischer, K. Goldmann, G. Le Provost, J. Overmann, L. Ruess, I. Schöning, S. Seibold, J. Sikorski, M. Neyret

## Abstract

Ecosystems worldwide face threats related to human-driven degradation, climate change, and biodiversity loss. Addressing these challenges requires management strategies that combine biodiversity conservation with climate change mitigation. Here, we aimed to identify local-scale management actions that promote biodiversity at multiple trophic levels while also promoting carbon storage and sequestration. We combined data on the diversity of nine taxonomic groups (plants, birds, moths, Mollusca, soil fungi, active soil bacteria, Cercozoan and Endomyxan soil protists, Oomycotan soil protists, and nematodes), with above- and belowground carbon storage in 150 temperate forest plots in three regions of Germany. These were dominated by European beech, pine, spruce and oak. We investigated the relationships between multiple forest structure and management variables, biodiversity and carbon storage and sequestration in forest plots with different management types. Carbon storage was 32% higher in uneven-aged than even-aged forests and increased with mean tree diameter, while carbon sequestration in trees was 15% higher in even-aged than uneven-aged stands. Mean tree diameter was positively related to overall biodiversity, especially bird species richness and the richness of forest specialist birds. Oak and beech-dominated stands harboured higher biodiversity than stands dominated by conifers (especially Pine). One exception to this was the richness of plant species and forest specialist plants, which were highest in spruce plantations. Surprisingly, deadwood input did not significantly affect the diversity of any taxonomic group in this study. By showing that older forests with a high proportion of uneven-aged broad-leafed trees, or oak-dominated forests, can promote both biodiversity and carbon storage, our results could help inform sustainable local-scale forest management in Central Europe that promotes both biodiversity conservation and carbon storage. These findings can form the basis of further larger-scale studies investigating such relations at larger spatial scales to inform landscape-level recommendations for sustainable multifunctional forest management.

## Introduction

Growing concerns on the repercussions of climate change and biodiversity loss on human well-being have led to increasing interest in ecosystem management strategies that tackle both threats (Millennium Ecosystem Assessment, 2005; Pettorelli et al., 2021; Turney et al., 2020; United Nations, 2021). These two challenges are often treated separately, but they share multiple drivers (Seddon et al., 2019) and climate change has also become a key driver of biodiversity loss (Lister & Garcia, 2018; Müller et al., 2023). As a result, the global community is under increasing pressure to address both crises simultaneously (Corlett, 2020; IPBES, 2019; Pörtner et al., 2021).

Forests, as one of the Earth’s primary carbon sinks (European Environment Agency, 2016) and home to high biodiversity (de Lima et al., 2020; Leuschner & Homeier, 2022) are often at the core of climate and biodiversity protection policies. Forests are estimated to store about 45 % of organic carbon worldwide (Bonan, 2008) >80% of aboveground carbon, and >70 % of soil organic carbon to a depth of one metre (Birdsey et al., 1993; D. D. Richter et al., 1999; Six et al., 2002; Wellbrock et al., 2017). Many forest ecosystems are recognized as biodiversity hotspots (Soto-Navarro et al., 2020), hosting most of the Earth’s terrestrial species (e.g. 80% of amphibian species, 75% of birds and 68% of mammals (FAO and UNEP, 2020)). Despite a high overall importance of forests for both biodiversity and carbon sequestration, high levels of both do not always correspond (Sabatini et al., 2019). In managed forests, stand-level forest management often focuses on narrow objectives like timber production (Simons et al., 2021), which shapes the vegetation structure and species composition of the stand (Felipe-Lucia et al., 2018). This impacts biodiversity (Brockerhoff et al., 2008; Brunet et al., 2010; Penone et al., 2019) and the ecosystem’s capacity to store carbon in soils and vegetation (Asbeck et al., 2021; Huston & Marland, 2003; Mayer et al., 2020).

In Germany, forests cover 32% of the land and provide employment for more than 1.1 million people (DFWR, 2022). Almost half (∼ 48%) of the forest area is privately owned. The other half is owned by the federal states (∼29 %), communities (∼ 19%) and the federal government with ∼ 4% (BMEL, 2018). German forests have been shaped by a long history of forest management (DFWR, 2022; Gossner, 2013; Grove, 2002). Without human intervention, it is estimated that 92% of German forest area would be dominated by European beech (*Fagus sylvatica*) and oak (*Quercus petraea* and *Quercus robur.*) (Bohn et al., 2007; DFWR, 2022). However, from the 18^th^ up to the late 20^th^ century, conifer monocultures of Scots pine (*Pinus sylvestris*) and Norway spruce (*Picea abies*) were strongly promoted in Central Europe (Heinrichs et al., 2019; Knoke et al., 2008), resulting in the current national forest composition dominated by four genera: spruce, pine, beech, and oak (BMEL, 2018). Today, the German forestry system is moving away from a production-focused forestry, towards a multi-objective management system. Current guidelines aim to develop ‘ecologically and economically valuable forests’ through ‘close-to-nature’ forest management practices. This includes favouring structurally diverse and mixed stands and long management cycles (DFWR, 2022) and promoting and retaining habitat trees (Dörfler et al., 2020), with expectations that this will promote biodiversity. For instance, greater habitat complexity in uneven-aged and mixed forests promotes biodiversity at the stand scale (Penone et al., 2019), while large trees provide numerous microhabitats (Michel & Winter, 2009; Vuidot et al., 2011; Winter & Möller, 2008). Deadwood left in the stand is also thought to serve as a habitat and nutrition source for a wide range of species (Dittrich et al., 2014; Löfroth et al., 2023; Oettel et al., 2020; Sandström et al., 2019; Scott & Brown, 2008; Siitonen, 2001; Stokland et al., 2012). The impact of the dominant tree species, though, varies across taxa (Edelmann et al., 2022; Leidinger et al., 2021), but in general broad-leafed forests seem to be preferred by more species overall (e.g. Abele et al., 2014; Charbonnier et al., 2016; Russ & Montgomery, 2002). This knowledge has accumulated in a piecemeal fashion, with studies focusing on either a few taxonomic groups (e.g. Leidinger et al., 2020) or single management variables (Sandström et al., 2019; Schulze, 2018). As a result, trade-offs and synergies between different taxa across multiple forest types have not been quantitatively assessed. A more complete assessment of how these management practices affect the diversity of multiple taxa, as well as forest potential for climate mitigation, could help assess the suitability of current management guidelines and support the sustainable use and conservation of German forests.

In this study, we investigate how forest structure affects synergies and trade-offs between biodiversity and carbon storage and sequestration in Central European forests. We combine data on the alpha diversity of above- and below-ground taxa from nine taxonomic groups, carbon storage and sequestration, and forest structure and management variables collected in 150 forest plots with different management types across Germany. We created indices combining either carbon- or biodiversity-related variables (Biodiversity and Carbon indices) to summarise the impacts of forest management variables on each of these two dimensions, as well as their joint response. We then assessed trade-offs and synergies between these aggregate metrics with linear (or generalized linear) models. We hypothesized that (1) carbon storage in trees is higher in older forests, (2) deadwood retention contributes positively to soil carbon storage, (3) biodiversity is higher in uneven-aged forests with older trees with abundant deadwood, and in mixed or broad-leafed forests rather than coniferous forests, and (4) both Biodiversity and Carbon indices increase with thicker tree diameter and decrease in coniferous stands. If these hypotheses are supported, it would indicate that local-level forest management that lengthens rotation cycles and promotes structural diversity might concurrently promote carbon storage and biodiversity conservation at the stand level.

## Methods

### Study sites and design

This study is part of the Biodiversity Exploratories project (<u>biodiversity-exploratories.de</u>), a large-scale and long-term project located in three regions of Germany: Schwäbische Alb in the south-west, Hainich-Dün in the centre, and Schorfheide-Chorin in the north-east. Each Exploratory comprises 50 forest plots (100 m × 100 m) selected to span the typical range of local tree species composition and management types. The regions were selected to be typical of the major different climate and geology types within Germany (except for the Alps and riparian ecosystems), and are also broadly representative of the most common forest types of Central Europe (Fischer et al., 2010). Further details on methods and data acquisition can be found in Table S1.

### Data acquisition

All data preparation and analysis was conducted with R version 4.3.1 (R Core Team, 2023).

### Forest management and structure data

Forest features were measured during two comprehensive forest inventories between 2008 and 2010 and 2014-2018, respectively. In each plot, all trees with a diameter at breast height (DBH) > 7 cm were surveyed. We focused on five forest management and structure variables (see Table S1 for details): total deadwood input per year (supply rate of deadwood to consumers), mean tree diameter at breast height (DBH), the degree of forest mixture (1 minus the proportion of the most abundant genus based on crown projection area), the identity of the dominant genera (pine, spruce, oak and pine), and the management type (uneven aged and even aged). Uneven-aged stands included both unmanaged plots and those based on a single-tree selection system. When multiple layers were present, they were combined for the calculation of the forest structure variables.

### Carbon storage

We calculated two measures of forests’ capability to store carbon: *C storage* was estimated from carbon stocks in soil and in the tree biomass. *C sequestration* was estimated from the annual increment of C in the trees. C stocks in deadwood and C fluxes from soils and vegetation were not considered as these are difficult to estimate accurately over meaningful timescales across many plots. Soil organic carbon storage was measured in 2014 in the topsoil (0-10 cm depth) using the dry combustion analysis. We calculated the tree carbon storage from standing wood volume, measured between 2014 and 2018. To obtain the aboveground *C storage* for each plot, we summed up the C storage for all tree species recorded in the plot. We calculated the *C sequestration* by using the annual wood increment measured between two second inventories conducted in 2008 – 2011 and 2015 – 2016, respectively. The total volume and volume increment was then multiplied by the plot’s average wood density, then multiplied by 0.5 to represent the proportion of mass that is carbon, and summed up per plot to obtain plot-level carbon sequestration. The average wood density used in this approach was calculated from the percentage of basal area occupied by each species in the plot and multiplied by species-specific wood densities from Vries et al. (2003).

### Biodiversity

We considered nine taxonomic groups; plants, birds, moths, Mollusca, soil fungi, the active fraction of soil bacteria, soil protists of Cercozoa and Endomyxa, soil protists of Oomycota, and nematodes, to represent a complete picture of the taxonomic diversity of below- and aboveground groups. The diversity of individual groups was measured at different time points during the 2015-2018 sampling period (see details in supplementary material Table S1). When multiple sampling years were available, we used the most recent or most complete data. We calculated species richness for each individual group using the R package ‘vegan’ (Oksanen et al., 2020).

We used two main indicators for biodiversity. First, we calculated the overall ecosystem richness considering all taxonomic groups (i.e., ‘multidiversity’ *sensu* Allan et al., 2014). Multidiversity is calculated as the average scaled species richness per taxonomic group, where the species richness of each group is scaled to its maximum across all plots (Allan et al. 2014). An advantage of the multidiversity metric over total species richness is the equal weighting of the species, thus preventing the measure from being driven by species-rich groups. As a result, plot-level multidiversity values vary between 0 (all groups simultaneously have their lowest observed richness) and 1 (all groups simultaneously have their highest observed richness) (Allan et al., 2014).

Second, we selected indicators of biodiversity representing high conservation value. We calculated the species richness of red-listed bird species in Germany (including category 1 (Critically Endangered), 2 (Endangered) and 3 (Vulnerable) (Grüneberg et al., 2016)) and the richness of birds and plant forest specialists (Table S2). Plant forest specialists were classified as plants species only found in forests, including open areas in forests (Schmidt et al. 2011). For birds, we used the European forest bird specialists of the list by Gregory et al. (2007). All species considered high conservation value are listed in Table S2.

### Correction for environmental covariates

The study regions differ greatly in climatic and geological conditions and the effect of these on biodiversity and carbon storage could mask that of local forest management. To assess the effect of our focus variables independently of environmental covariates, we first corrected for environmental covariates. To do so, we selected five environmental covariates that represent soil, climatic and topographic conditions and with relatively low correlation: soil pH, mean annual temperature, soil depth, proportion of clay in the soil, and the Topographic Wetness Index (Moeslund et al., 2013). We fitted linear models for each response variable with all environmental covariates as well as the region as explanatory variables. We then extracted the residuals from each model. To maintain the relative ratios of the multiple variables, we added the mean value of each response variable to these residuals; this was especially useful for carbon storage in trees or soil, which are measured on very different scales. These adjusted values were then used in all further analyses. To ensure normal error distributions and a homogeneous variance, a square root transformation was applied to deadwood input.

### Statistical analysis

Since the main objective of the study was to identify the conditions that simultaneously maximise biodiversity and carbon storage/sequestration, we created indices combining multiple carbon and/or biodiversity variables (Figure 1). At each aggregation step, variables were scaled between 0 and 1 to ensure equal weighting of all response variables (Manning et al. 2018), hence all indices ranged from 0 to 1. The ‘*Carbon index*’ was measured as the average of tree and soil carbon storage and tree carbon sequestration, equally weighted. The ‘*Biodiversity index*’ was calculated from the multidiversity measure (Allan et al. 2014) and ‘Conservation species index’, itself including the species richness with conservation value. Finally, the ‘*Combined index*’ was calculated as the average of carbon and the biodiversity indices (Figure 1). There were a few missing values: 11 for Mollusca, 3 for Cercozoa and Endomyxa Protists and moths, 1 for birds, oomycote and Nematodes, and 14 for deadwood input. Because deadwood input was an explanatory variable in all models, we filled missing values with the average of deadwood input in all plots. Other NAs were not imputed.

**Figure 1:**
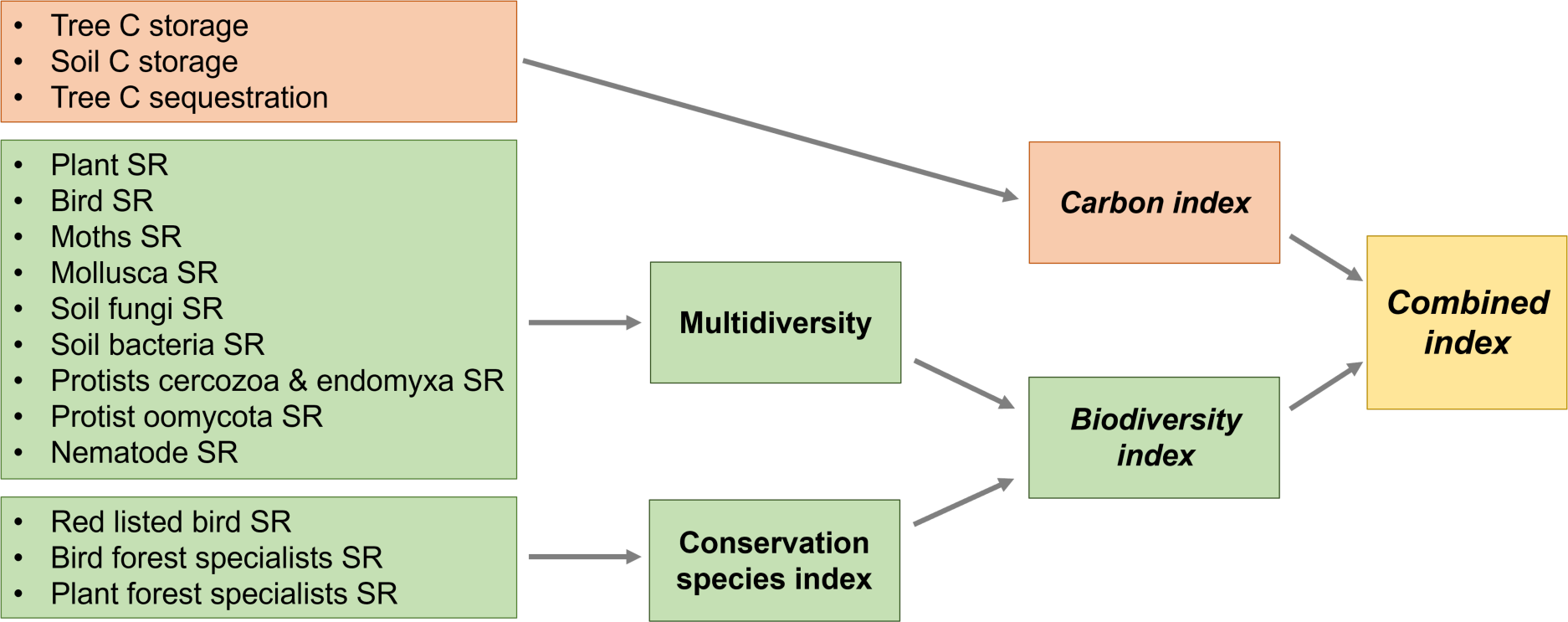
Schematic illustration showing the calculation of the Combined index. All the variables and intermediate indices were adjusted and scaled between 0 and 1. Multidiversity was calculated according to Allan et al. (2014). Grey arrows indicate scaling and averaging of variables into the next level of aggregation. SR = species richness.

Finally, we fitted linear models (function: lm; package: *stats*) between each response variable (richness, carbon variables, or indices) and above-mentioned forest structure variables as explanatory variables. The exception was red-listed bird species, which were fitted using a Poisson glm. Model tests were conducted using variance analysis (ANOVA). Considering that not all variable combinations were available (e.g. no uneven-aged pine- or spruce-dominated stands) we did not investigate the interactions between the different variables. Model comparisons were done using the *parameters* package and forest plots using the *sjPlot* package.

## Results

### Management and forest structure variables

European beech was dominant in 70% of the plots (105 plots), followed by Scots pine (∼13%; 19 plots), and Norway spruce (∼11%; 17 plots). The two oak species were dominant in nine plots (*Quercus robur* and *Quercus petraea,* ∼6%). All spruce- and pine-dominated plots were managed as even-aged forests. This was also the dominant management type in plots dominated by beech (∼65%: 68 out of 105 plots) and oak (∼88 %: 8 out of 9 plots). Pure stands (91 plots) were more common than mixed ones (59 plots), except oak stands which were more mixed (6 stands) than pure (3) (Fig. S1).

Forests dominated by oak and spruce had a higher mean DBH (mean 35.1 cm ± sd 8.5 and 31.8 cm ± 6.4 respectively) than forests dominated by beech and pine (27.5 cm ± 11.9 and 27.4 cm ± 8.4 respectively). Mean DBH was also higher in uneven-aged than even-aged forests (32.3 cm ± 10.4 and 27.1 cm ± 10.9 respectively, p = 0.01) and in pure than mixed forests (31.2 cm ± 10.6 and 24.1 cm ± 10.2, p ≤ 0.001). This shows that mean DBH, dominant genera and management type were not fully independent from each other.

### Forest structure and management for a high *Carbon index*

Overall, the *Carbon index* ranged between 0.29 and 0.84 and tended to be higher in stands with large mean DBH and lower in pine-dominated stands (Fig. 2, Table 1). Soil carbon did not respond to any of the management variables. The component variables of the *Carbon index* differed in their respective responses to the explanatory variables. Tree carbon storage increased with mean DBH (standardised effect size: 0.65 ± 0.06 p < 0.001), as expected from its calculation from tree volume. Tree carbon storage was on average 32.13% higher in uneven-aged than even-aged stands (0.26 ± 0.06, p < 0.001). High tree carbon storage was also associated with high deadwood input (0.13 ± 0.05 p = 0.009), likely due to stands with larger, older trees also having higher tree senescence. Finally, tree carbon storage was 35% lower in pine-dominated stands compared to beech stands (−0.73 ± 0.16, p < 0.001). Conversely, carbon sequestration was 15.07% lower in uneven-aged stands (−0.37 ± 0.09 p < 0.001) and tended to decrease with mean DBH (−0.13 ± 0.09, p = 0.1, Fig. 2). These results support our hypothesis 1, since carbon storage in trees was higher in forests with a higher mean DBH, and rejects hypothesis 2, that deadwood retention contributes to soil carbon storage, since we could not identify a significant relation between any management variable and soil carbon storage.

**Figure 2:**
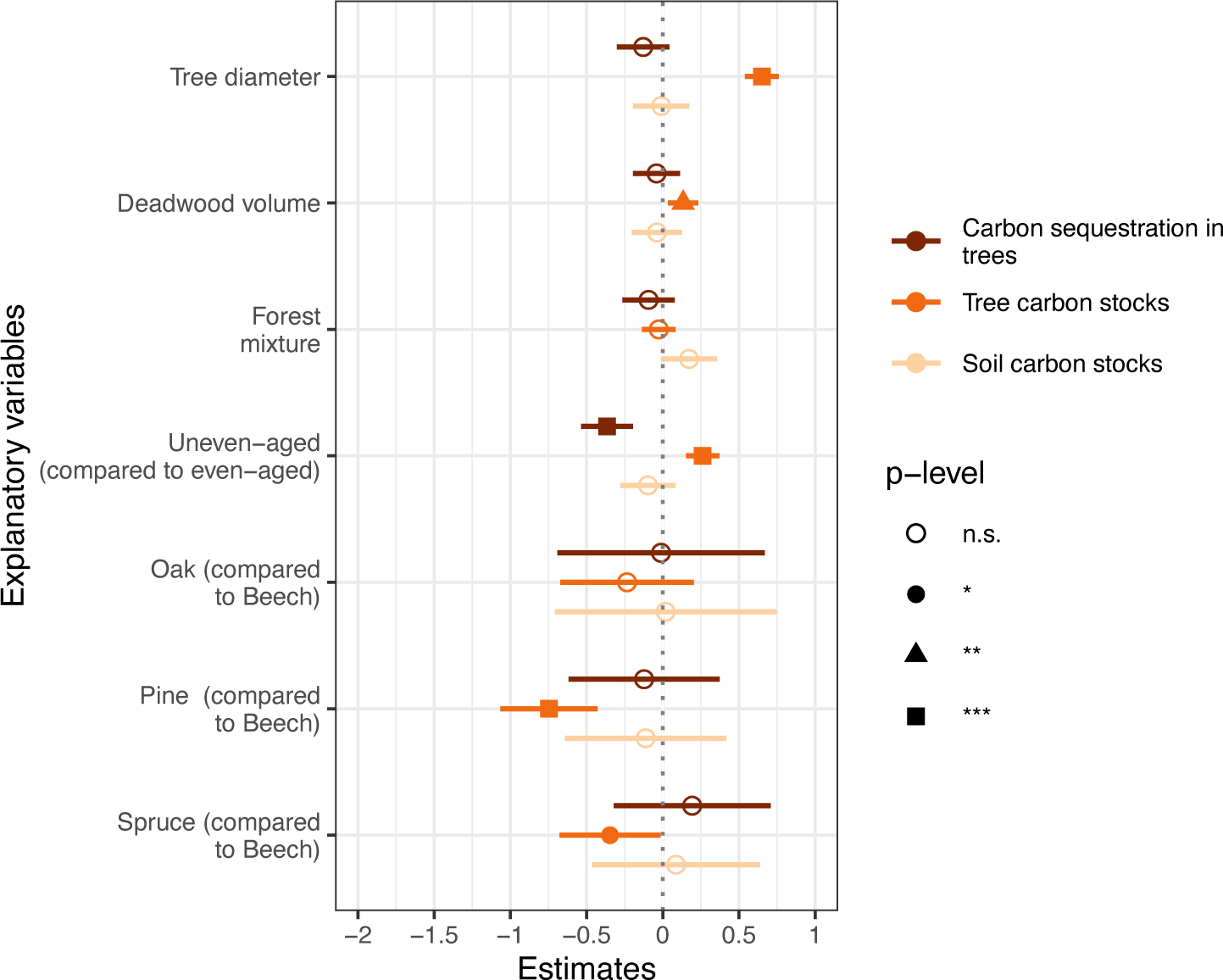
Effect of forest properties on carbon stocks and sequestration. Plots show standardized effect sizes along with 95% confidence interval estimated for the selected management variables affecting carbon stocks in trees and soils and carbon sequestration in trees. For dominant genus, the results are shown in comparison with the reference genus (beech, the most abundant genus). For stand age structure, the results are shown for uneven-aged stands in comparison to even-aged stands. SR = species richness. * p < 0.05, ** p < 0.01, *** p < 0.001, n.s. = non-significant.

**Table 1:** Linear model results for the effect of management on the response variables showed as F value (for all groups except red listed birds for which deviance is shown instead). Significance is shown as * 0.05 < p ≤ 0.01; ** 0.01 < p ≤ 0.001; *** p < 0.001, and • reflects marginal effects (0.1 < p ≤ 0.05). The degrees of freedom (df) for each effect are shown for each explanatory variable. Response variables are tree carbon, soil carbon, carbon sequestration, multidiversity, plant specialists, bird specialists, red listed birds and the three indices.

| Explanatory variables | Df | Tree C storag e | Soil C storage (kg.m<sup>-2</sup>) | Tree C sequestr ation | Multi- diversity | Plant specialis ts | Bird special ists | Red listed birds | Carbon Index | Biodiver sity Index | Combin ed Index |
| --- | --- | --- | --- | --- | --- | --- | --- | --- | --- | --- | --- |
| Mean DBH | 1 | 191.3 | 0.8 | **4.3\*** | **19.6\*\*\*** | 2.2 | **17.1\*\*\*** | 6.9 | **11.8\*\*\*** | **19.5\*\*\*** | **32.6\*\*\*** |
| Deadwood input | 1 | 6.8 | 0.4 | 0.5 | 0.9 | 1.0 | 1.4 | 0 | 0 | 0.9 | 0.42 |
| Genus | 3 | **7.5\*\*\*** | 0.1 | 0.3 | **3.3\*** | **8.3\*\*\*** | 2.0 | **7.8.** | 2.0 | **3.3\*\*** | **4.8 \*\*** |
| Mixture | 1 | 0 | 4.6* | 0.6 | **2.8 .** | **3.5 .** | **8.4\*\*** | 0.4 | 0.6 | **2.8 .** | **3.1 .** |
| Management type (even/uneven-aged) | 1 | **49.7\*\*\*** | 1.3 | **22.3\*\*\*** | 0.3 | 16.6*** | 0.4 | **8.7\*** | 2.0 | 0.3 | 0.6 |

### Forest structure and management for a high *Biodiversity index*

The diversity of individual taxa showed contrasting responses to forest structure and management variables, but overall increased with mean DBH and decreased in pine-dominated stands (Fig. 3). Individual responses of the species richness of all considered groups can be found in Fig. 3, Table 1 and Table S3. There was strong evidence that mean DBH positively affected bird (0.28 ± 0.09 p = 0.002) and forest bird specialist species richness (0.38 ± 0.09, p < 0.001). It also positively affected the richness of cercozoan and endomyxan (0.21 ± 0.09 p = 0.016) as well as Oomycotan protists (0.19 ± 0.09 p = 0.041). Forest mixture had a more variable impact, affecting positively the richness of soil fungi (0.29 ± 0.09 p = 0.002), and bird specialists (0.18 ± 0.09 p = 0.038), but negatively the richness of plant forest specialists (−0.16 ± 0.08, p = 0.05). Uneven-aged stands had lower plant (−0.19 ± 0.08 p = 0.015), forest plant specialist (−0.23 ± 0.08 p = 0.005) and nematode (0.18 ± 0.09 p = 0.045) richness, but higher red-listed bird richness (0.29 ± 0.12 p = 0.014) than even-aged stands.

**Figure 3:**
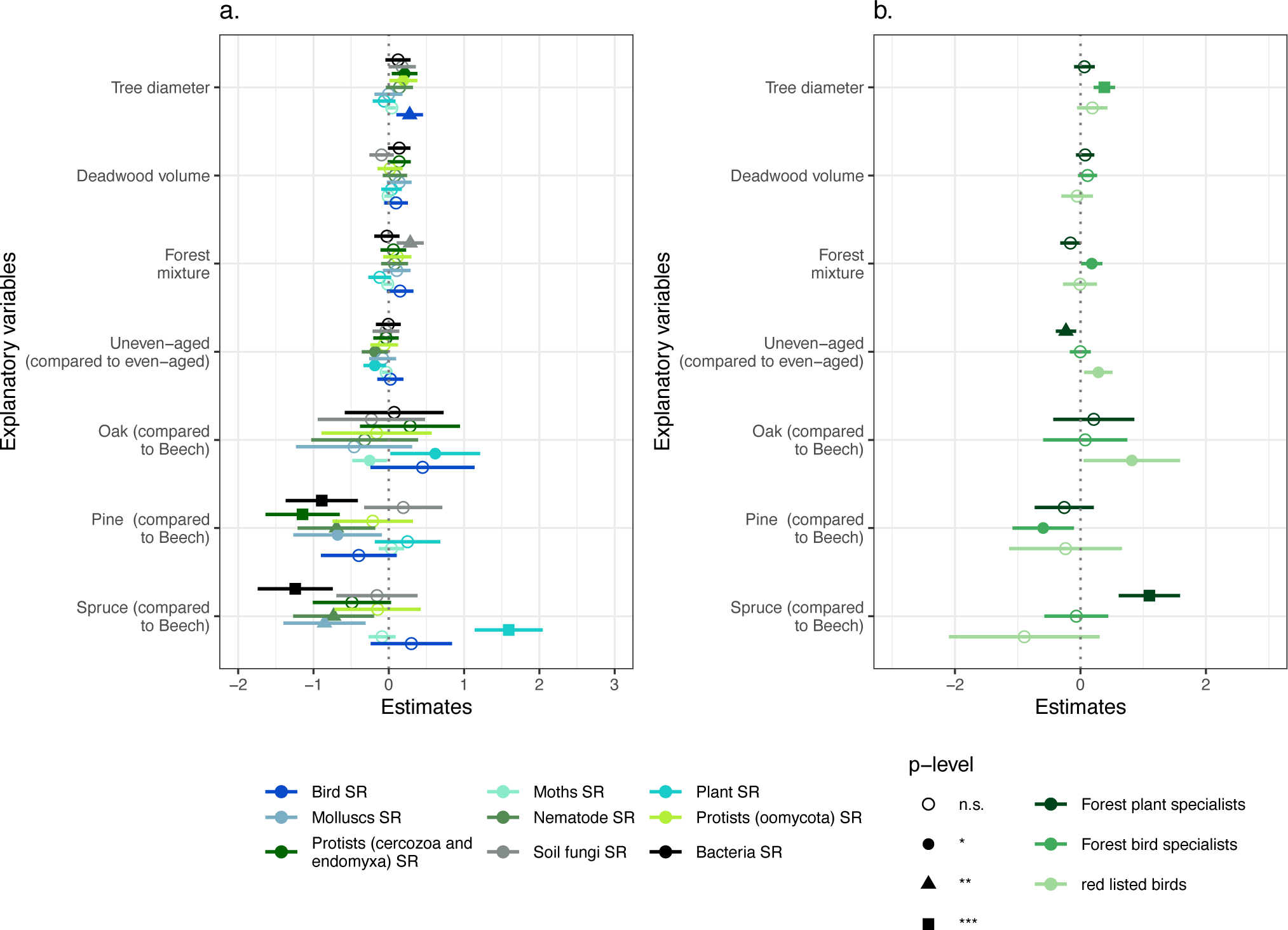
Effect of forest stand properties on biodiversity measures. Plot show standardized effect sizes along with 95% confidence interval estimated for the selected management variables affecting a) species richness of individual groups and b) species richness of groups with specific conservation value. For dominant genus, the the results are shown in comparison with the reference genus (beech, the most abundant genus). For stand age structure, the results are shown for uneven-aged stands in comparison to even-aged stands. SR = species richness. * p < 0.05, ** p < 0.01, *** p < 0.001, n.s. = non-significant.

There were also important but contrasting effects of the dominant tree genus on different taxonomic groups. Oak stands promoted higher plant (0.62 ± 0.30 p = 0.043) and red-listed bird richness (0.82 ± 0.39 p = 0.036) than beech stands, but lower moth richness (−0.25 ± 0.12 p = 0.031). Spruce stands had lower bacteria (−1.24 ± 0.25 p < 0.001), nematode (−0.73 ± 0.27 p = 0.008) and mollusc (−0.85 ± 0.28 p = 0.002) richness, but higher plant (1.59 ± 0.23 p < 0.001) and forest plant specialist richness (1.10 ± 0.25 p < 0.001) compared to beech-dominated stands. Pine stands had a strongly negative impact on biodiversity compared to beech stands, with negative impacts on the richness of bacteria (−0.89 ± 0.24 p < 0.001), protists (−1.14 ± 0.25 p < 0.001), nematodes (−0.69 ± 0.26 p = 0.009) molluscs (−0.68 ± 0.30 p = 0.024) and bird forest specialists (−0.59 ± 0.25 p = 0.018).

Overall, multidiversity was highest when mean DBH was high (0.29 ± 0.09 p = 0.001), and lowest in pine-dominated stands (−0.69 ± 0.25 p = 0.007). The diversity of species of high conservation value (forest specialists and red listed birds) increased with mean DBH (0.29 ± 0.09 p = 0.001) and were higher in Oak-dominated stands (0.71 ± 0.34 p = 0.041). These results mostly confirm our hypothesis 3, although contrarily to our expectations deadwood input did not affect biodiversity.

### Forest structure and management for biodiversity conservation and climate mitigation

Our analysis revealed trade-offs between carbon storage (maximised in uneven-aged stands at high mean DBH) and sequestration (maximised in un-even-aged stands, at low mean DBH), and between some biodiversity groups. For instance, plant richness was the highest in spruce plantations where the diversity of most other groups was the lowest; and overall the Biodiversity index tended to be higher in oak stands, but this was not significant. when aggregated into the *Carbon* and *Biodiversity indices*, responses were relatively consistent, and the *Combined index* (ranging from 0.31 to 0.74) followed similar trends to its components: it increased with mean DBH (p < 0.001) and was lowest in pine-dominated stands (p < 0.001, Fig. 4; Table 1). This overall confirms our Hypothesis 4 by showing that the *Combined index* can be maximised by a higher mean DBH and stands with primarily broad-leafed trees.

**Figure 4:**
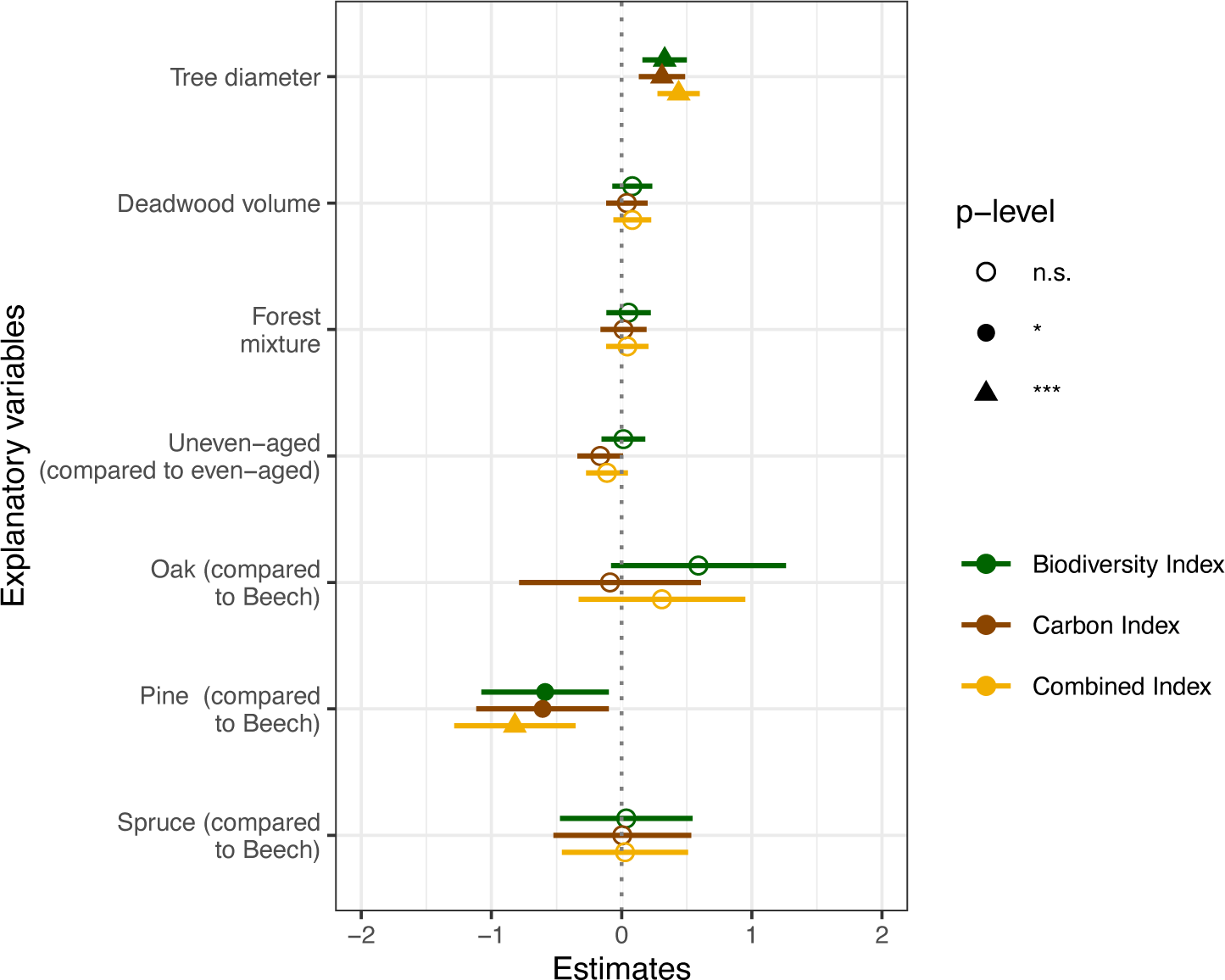
Standardized effect size along with 95% confidence interval estimated for the selected management variables affecting the *Biodiversity, Carbon and Combined indices*. For dominant genus, the results are shown in comparison with the reference genus (beech, the most abundant genus). For stand age structure, the results are shown for uneven-aged stands in comparison to even-aged stands. SR = species richness. * p < 0.05, ** p < 0.01, *** p < 0.001, n.s. = non-significant.

## Discussion

Our results show that carbon storage and biodiversity are typically simultaneously high in older broad-leaved-dominated stands with large diameter trees. However, beyond this simple conclusion are more nuanced relations between the forest, carbon and biodiversity properties. In the discussion we assess the relation of several forest properties to carbon and biodiversity, and discuss how these can be influenced by management.

### Forest structure and composition promoting carbon storage and biodiversity conservation

The dominant tree genus was an important driver that impacted 53% of the biodiversity and carbon variables. Differences were the strongest between pine- and beech-dominated stands, with pine-dominated stands having lower values of nine biodiversity indicators and lower tree carbon stocks. Spruce dominance was negatively related to three out of nine biodiversity indicators and positively related to two (plant species richness and forest plant specialist richness). These biodiversity patterns are consistent with previous studies showing that the plant species richness of German forests is high in coniferous stands (Boch et al., 2013; Budde et al., 2011) and can be explained by the fact that conifer canopies are more open than those of beech stands, allowing higher understorey light availability and more favourable microclimatic conditions (Dormann et al., 2020; Penone et al., 2019; Wagner et al., 2011). Oak-dominated stands had higher plant species richness and red listed bird richness but lower moth species richness than beech stands. Previous studies in Germany also found oak forests to be the most favourable to biodiversity (Carvalho-Santos et al., 2016; Müller et al., 2021). This can be explained by oaks being present in the study regions for centuries, and thus likely having a higher co-evolved diversity (Brändle & Brandl, 2001) than e.g. Scots pine which has been cultivated beyond its natural range, and for a much shorter time. Oaks also have higher microhabitat availability and dead branch accumulation than pines (Paillet et al., 2019). This might promote resource availability for insectivorous birds and provide more microhabitats for molluscs (Abele et al., 2014).

Of the forest structural properties we assessed, mean DBH and management type were strongly related to tree carbon storage, carbon sequestration rates and the biodiversity of multiple groups. Carbon sequestration was lower in forests with low mean DBH and in uneven-aged forests, likely due to slower growth rates in larger trees (Stephenson et al., 2014). This represents a classic trade-off in forest carbon management as stands with higher mean DBH have a higher tree carbon storage. The positive effect of DBH on many groups is likely due to the higher number of microhabitat types in larger trees (Michel & Winter, 2009; Vuidot et al., 2011; Winter & Möller, 2008), and an overall higher resource availability. Additionally, large trees are attractive for bird and arthropod cavity builders, as the high wood thickness of their cavities provides buffered microclimatic conditions for nests (Remm et al., 2006). Some of our results were more surprising, such as the positive association between mean DBH and protist diversity, which we could not explain, but which may be associated with the presence of a stable and long-lived habitat in which microbial diversity may accumulate. The benefits of habitat heterogeneity for biodiversity are also visible in the positive relationship between forest mixture and most of the biodiversity variables such as forest bird specialists and soil fungi (Leidinger et al., 2021).

Annual deadwood input was positively associated with tree carbon storage, probably because more deadwood is left in older stands with larger trees and because more trees are senescent in older forests. Deadwood is an important structural element in forests at it offers resources for biodiversity (Seibold et al., 2017) by storing large amounts of water, providing energy and nutrients to soil micro- and macro-organisms, and supplying habitats to xylobiontic species (Oettel et al., 2020; Scott & Brown, 2008). It has been estimated that, 20 - 25 % of all forest-dwelling species are dependent on deadwood (Siitonen, 2001). Yet, surprisingly, deadwood input did not significantly affect the diversity of any taxonomic group in our study. This was likely because (i) the taxa analysed did not include saproxylic and xylobiontic species, which are expected to most respond most strongly to deadwood (Sandström et al., 2019) and (ii) only few of the stands had a high rate of deadwood input (mean: 1.2 m^2^.ha^-1^.yr^-1^, max: 28.9 m^2^.ha^-1^.yr^-1^).

### Implications for German forest management

German forests are the product of a long history of forest management (DFWR, 2022; Gossner, 2013). In the last centuries, conifer monocultures were promoted in Central Europe (Heinrichs et al., 2019; Knoke et al., 2008; Penone et al., 2019), leading to the current national forest composition, with almost 75 % of the total forest area dominated by four genera: spruce (25 %), pine (23 %), beech (16 %), and oak 10 % (BMEL, 2018). Compared to this national average, in our study beech was overrepresented (70% stands), while spruce is underrepresented, but overall our study plots covered most of the main forest stand types found in Germany.

Current guidelines adopted in Germany aim to develop ‘ecologically and economically valuable forests’ through ‘close-to-nature’ forest management practices. These include the promotion of structurally diverse and mixed stands and long cycles (DFWR, 2022). As part of this, spruce and pine forests are being converted into mixed stands (Ammer, 2019; Ammer et al., 2008; Heinrichs et al., 2019; Knoke et al., 2008; von Lüpke et al., 2004), and broad-leafed tree cover has increased steadily (+7% between 2002 and 2012, BMEL, 2018). Our results show that these changes are likely to benefit both biodiversity and carbon storage through decreased coniferous (especially pine) cover and a switch from monocultures to mixed forests with larger resource heterogeneity (Heinrichs et al., 2019). Longer forest cycles and thus forest in late development stages, are also becoming more common (BMEL, 2018). This will benefit biodiversity and climate-friendly forestry since large diameter trees store more carbon than young trees (European Environment Agency, 2016), although the growth and thus carbon sequestration is overall lower in older trees (Johnson & Abrams, 2009, Meinzer et al. 2011). Overall, our results confirm the idea that young stands will allow for new carbon stocks to be sequestered, but that old broad-leafed stands should be kept as long as possible, to promote both carbon storage and biodiversity.

### Future directions

Our results confirm that the current management trends in German forestry should promote more biodiverse and climate-friendly forests at the local stand level. However, other elements need to be assessed for a more comprehensive understanding of the impact of forest management on wider scale biodiversity, other aspects of carbon cycling, and other ecosystem services. More specifically, young stands may become more prevalent in the future due to increasing rates of disturbances due to bark beetles, wind and drought (Seidl et al., 2014; Senf & Seidl, 2021). The above-described trade-off between carbon storage and sequestration means that these young stands will accumulate carbon rapidly but will take time to store significant amounts of carbon. Our results add to this by showing that these new stands may also take time to reach the high biodiversity values of current old forests - although they may support their own distinct biota. This highlights the importance of old forests, which can also act as biodiversity reservoirs from which species can colonise younger stands. The specific tree species (and associated management practices) being promoted will also influence the outcome, considering the role of the dominant tree genus on both biodiversity and carbon storage shown above (Felton et al., 2010).

In this study we considered the impact of stand management on just two aspects of forest carbon: tree and topsoil carbon storage. For a complete picture of climate mitigation, it is also important to account for carbon sequestration and storage in deeper soil layers and the rest of the wood production line (e.g. carbon release from use as firewood vs. long-term storage as timber, effects of thinning). Accounting for these processes would provide more precise estimates of the relative impacts of different forest management practices. The inclusion of other greenhouse gases would also provide a more complete assessment of climate impacts. Finally, we only assessed the impact of forest management on biodiversity and climate change mitigation. Yet, forests provide a wider range of ecosystem services such as the production of timber, the regulation of water and air quality and also have cultural and recreational value (Führer, 2000; Neyret et al., 2023). Different ecosystem services might be favoured by different forest types, and their consideration could highlight additional trade-offs and synergies (Felipe-Lucia et al. 2018) for local management recommendations. Promoting these different services might require maintaining landscape-level diversity in the type and management of forests, likely resulting in a higher overall forest management diversity due to diverse species requirements (Schall, Gossner, et al., 2018; Schall et al., 2020). While our results indicate optimal forest stand properties for maximizing local-scale biodiversity and carbon, we also recommend maintaining forest diversity and heterogeneity at the landscape level. This can help promote landscape-level multifunctionality, whereby different forest stands simultaneously provide biodiversity protection, climate mitigation options, economic benefits, as well as cultural values (van der Plas et al., 2016).

## Conclusion

Simultaneously promoting biodiversity protection and climate change mitigation is a key challenge of local-scale forest management. Here, we identified the forest management features that support these goals in Central European forests: large average tree diameter and dominance of species such as oak. This study highlights the need to pay special attention to old forests due to their importance for biodiversity and carbon storage; further research should however build on these results to assess their resilience to future climates as well as the role of forests with different compositions than those assessed here. As the demand for preserving both climate and biodiversity grows stronger, approaches such as that presented here can help support management decisions and forest management policies, and thus promote more sustainable and multifunctional forests.

## Acknowledgements

We thank the Exploratories managers: Konstans Wells, Swen Renner, Kirsten Reichel-Jung, Sonja Gockel, Kerstin Wiesner, Katrin Lorenzen, Andreas Hemp, Martin Gorke and all former managers for maintaining the plot and project infrastructure; Simone Pfeiffer, Maren Gleisberg and Christiane Fischer for central office support, Jens Nieschulze and Michael Owonibi for database management, and Eduard Linsenmair, Dominik Hessenmöller, Daniel Prati, François Buscot, Ernst-Detlef Schulze, Wolfgang W. Weisser and the late Elisabeth Kalko for establishing the Biodiversity Exploratories project. We also thank all data contributors.

The work has been funded by the DFG Priority Program 1374 “Infrastructure-Biodiversity-Exploratories”. Fieldwork permits were issued by the responsible state environmental offices of Baden-Württemberg, Thüringen, and Brandenburg.

## Data availability

This work is based on data elaborated by several projects of the Biodiversity Exploratories program (DFG Priority Program 1374). Most datasets are publicly available in the Biodiversity Exploratories Information System (http://doi.org/10.17616/R32P9Q). However, to give data owners and collectors time to perform their analysis the Biodiversity Exploratories’ data and publication policy includes by default an embargo period of three years from the end of data collection/data assembly which applies to the remaining datasets. These datasets will be made publicly available via the same data repository. All datasets are listed in Tables S1 and corresponding references.

### Appendix

**Table S1.**
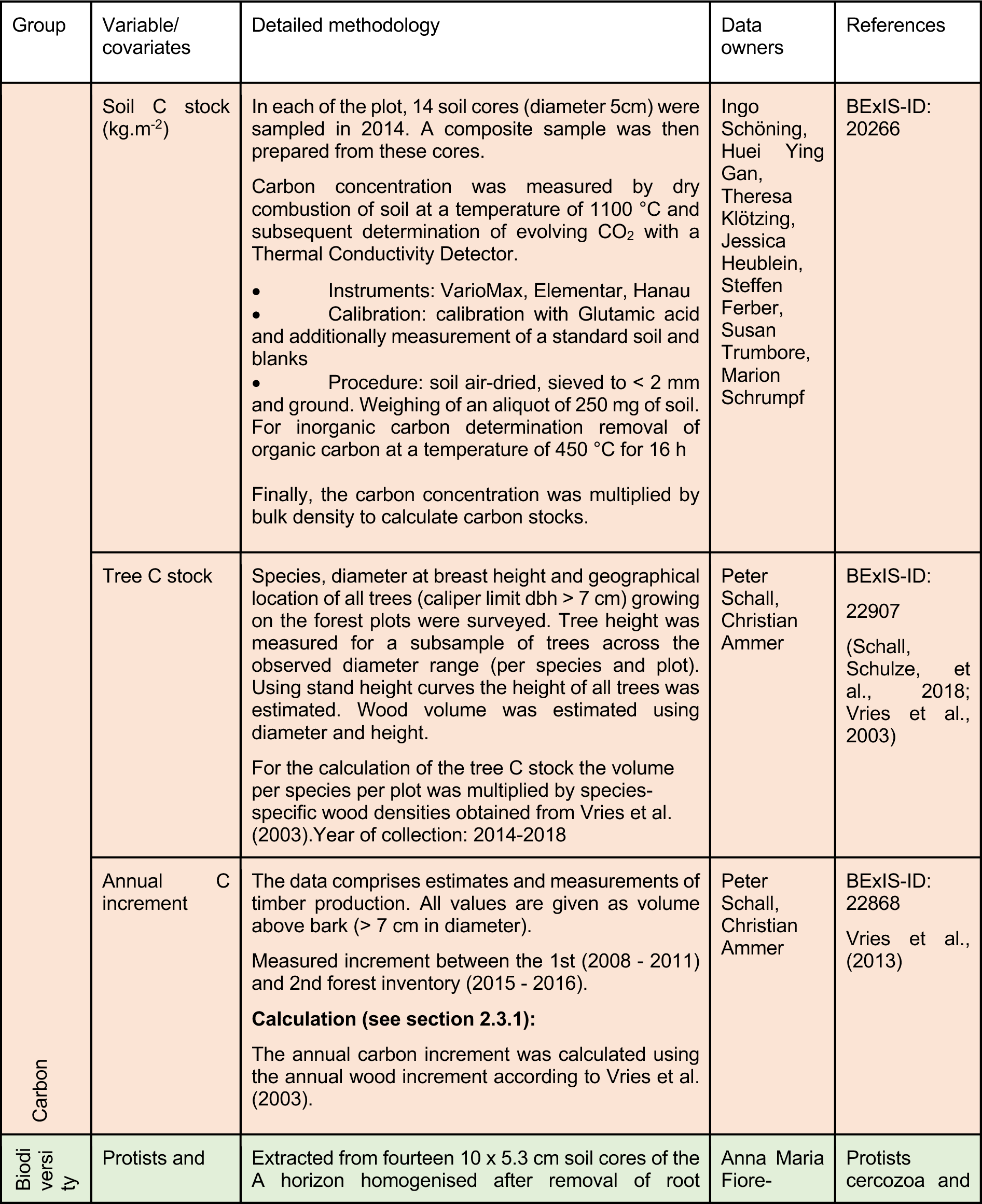
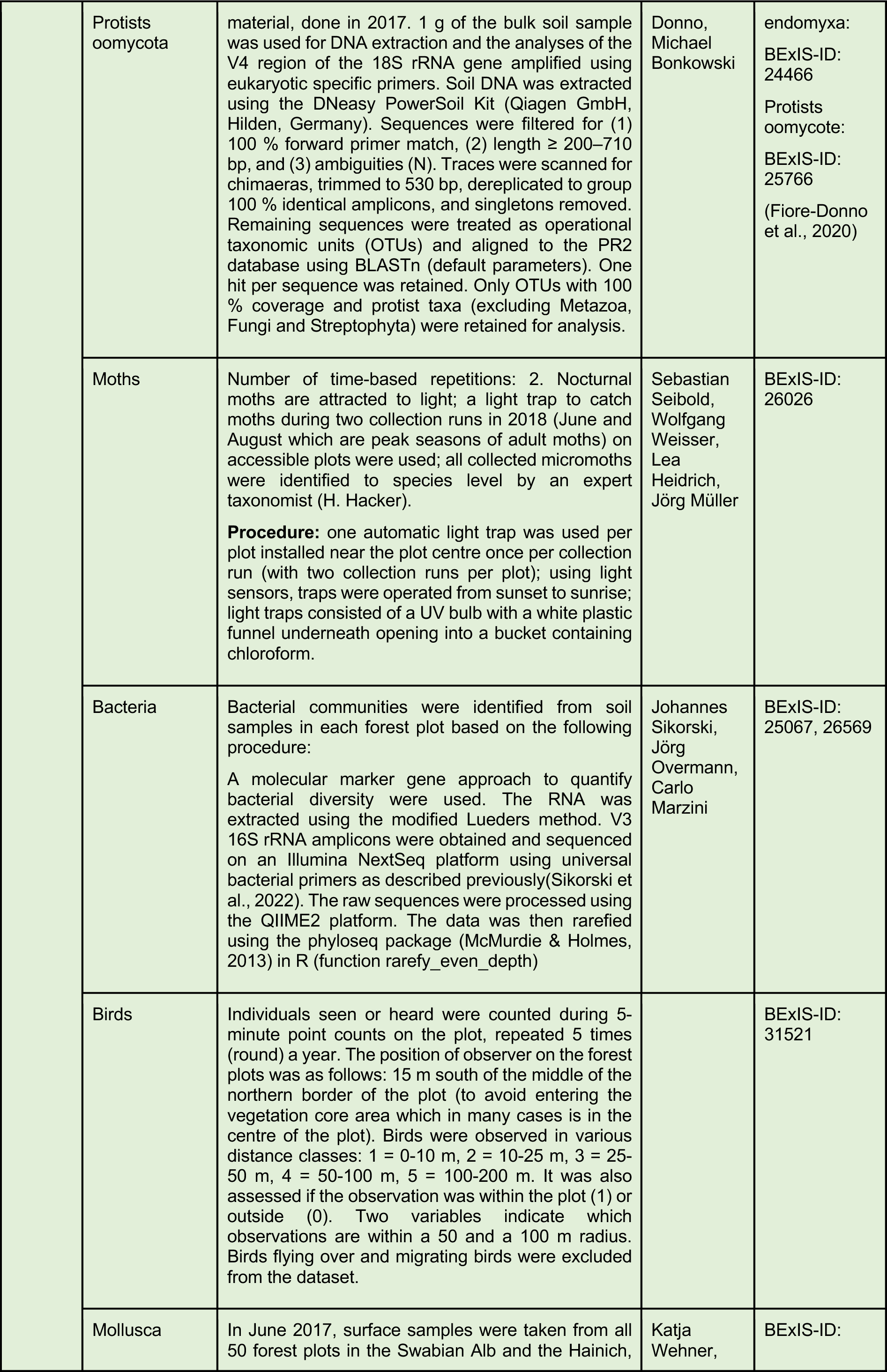
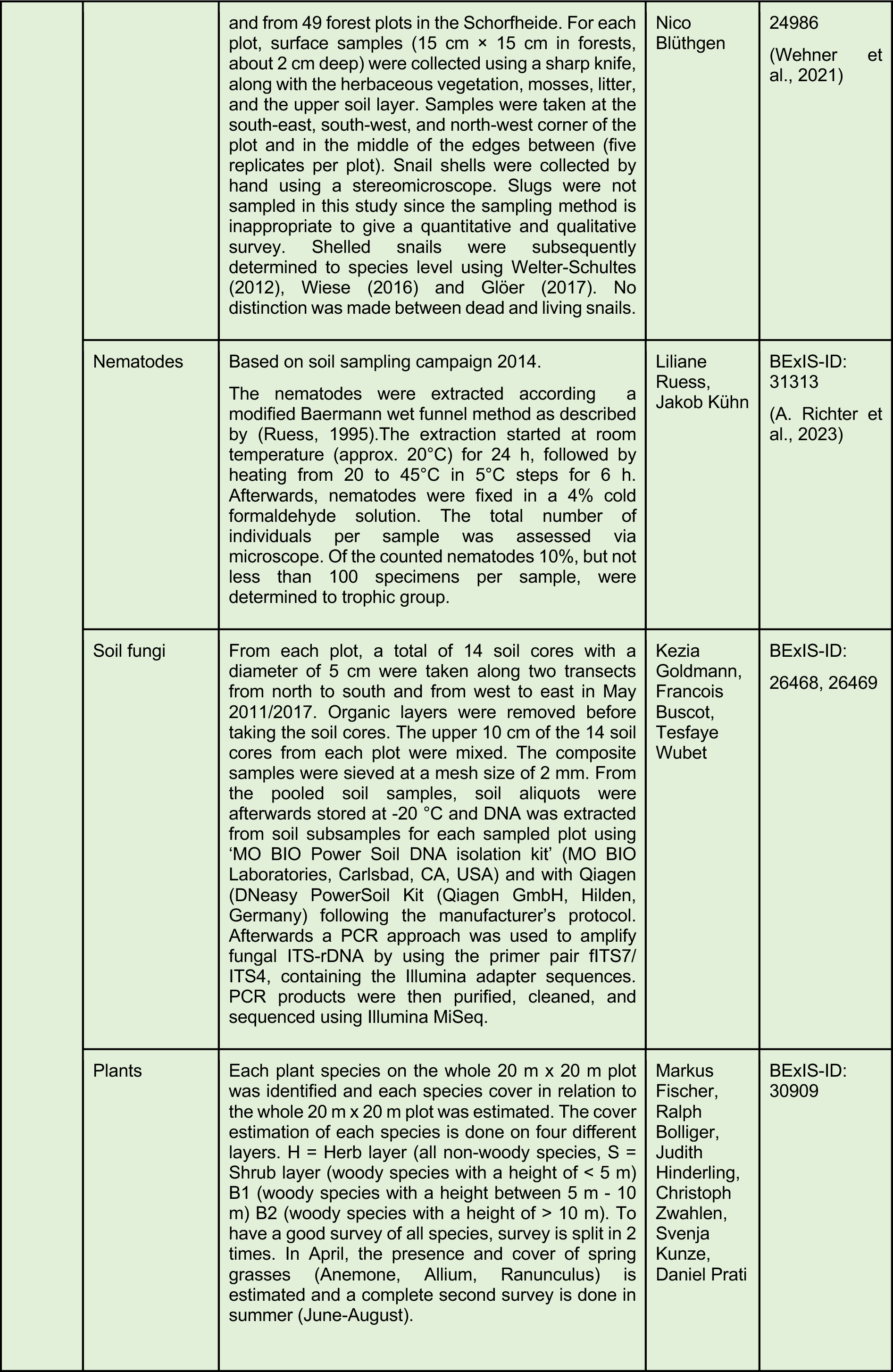
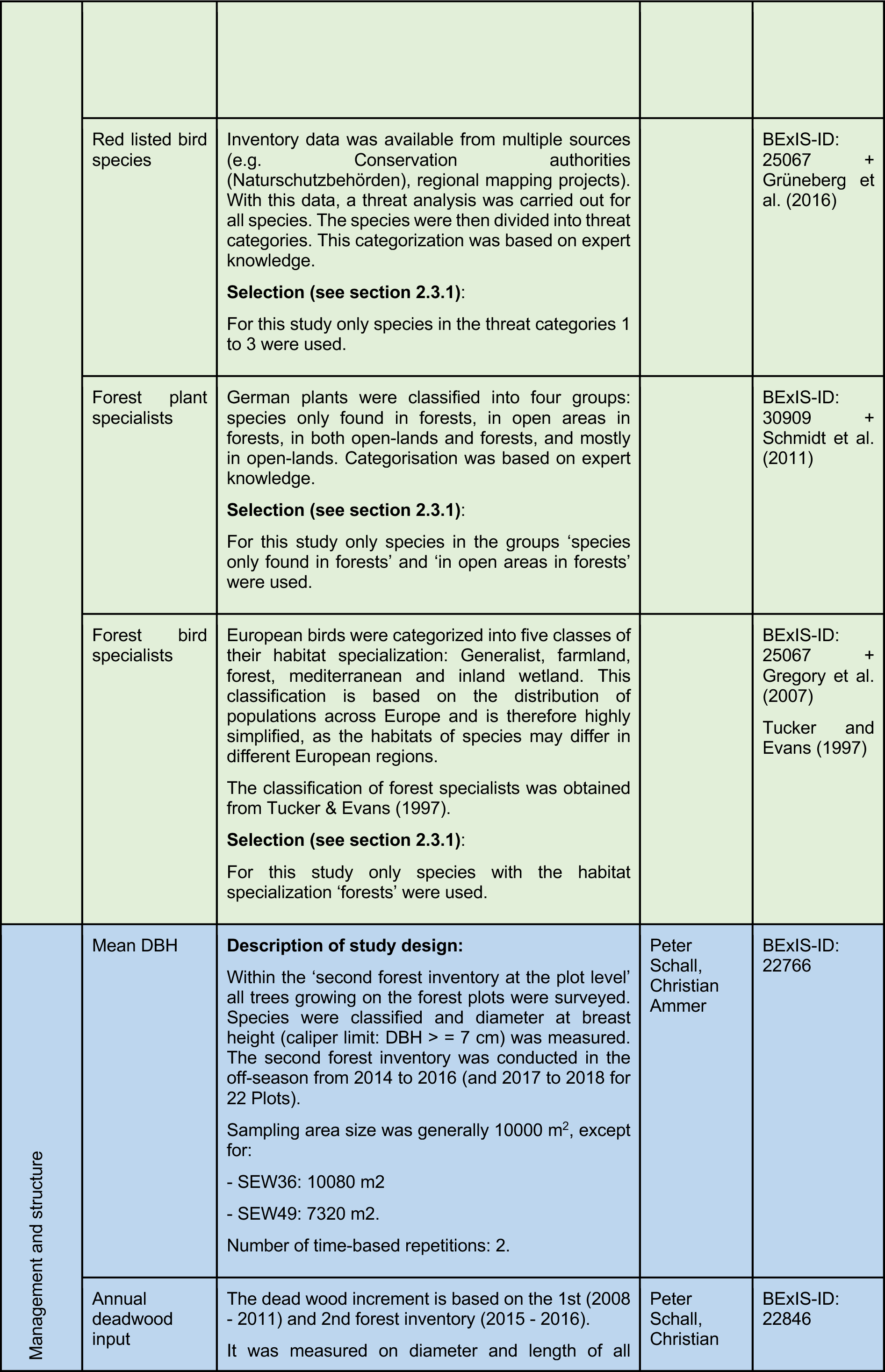
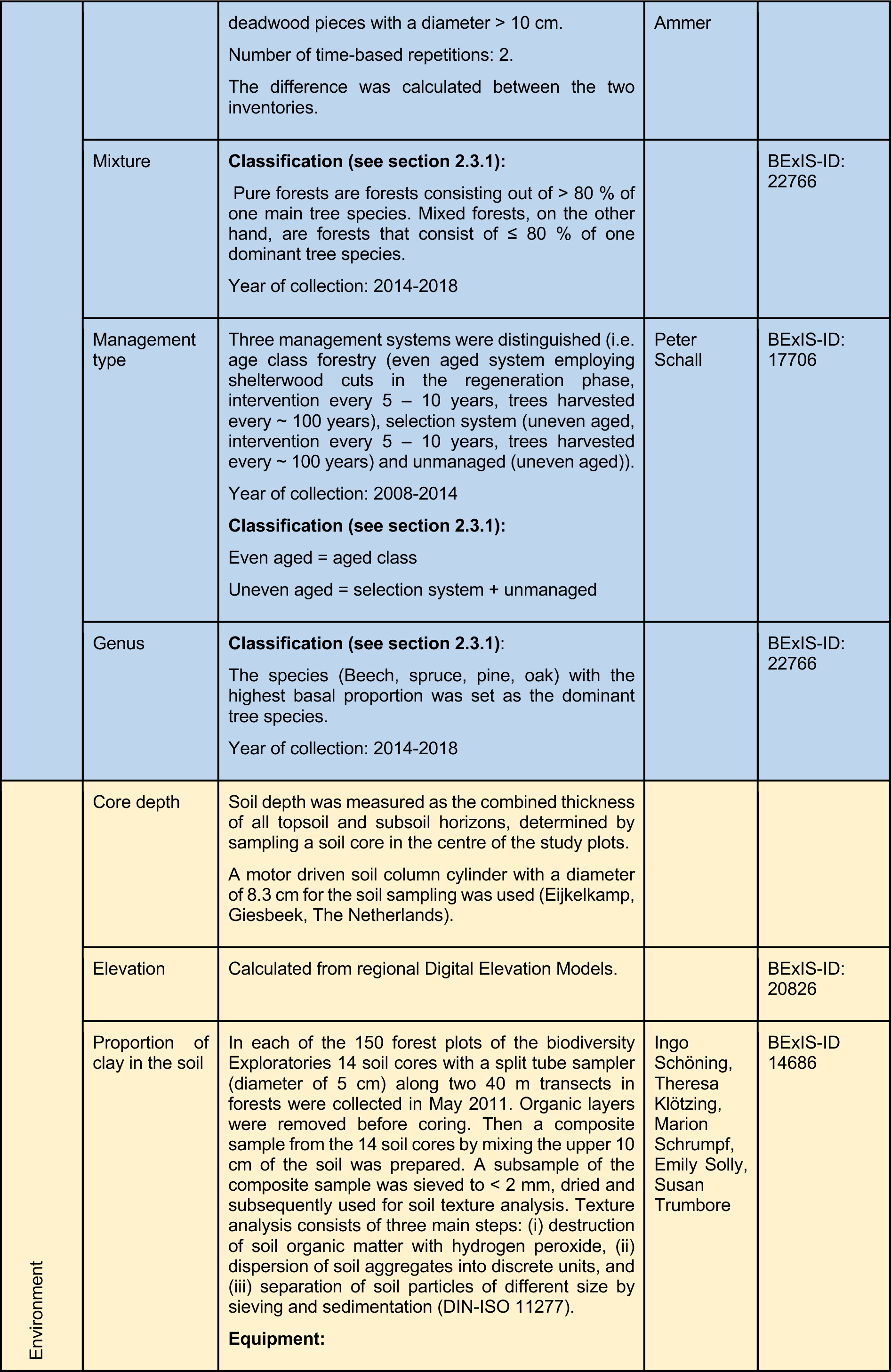
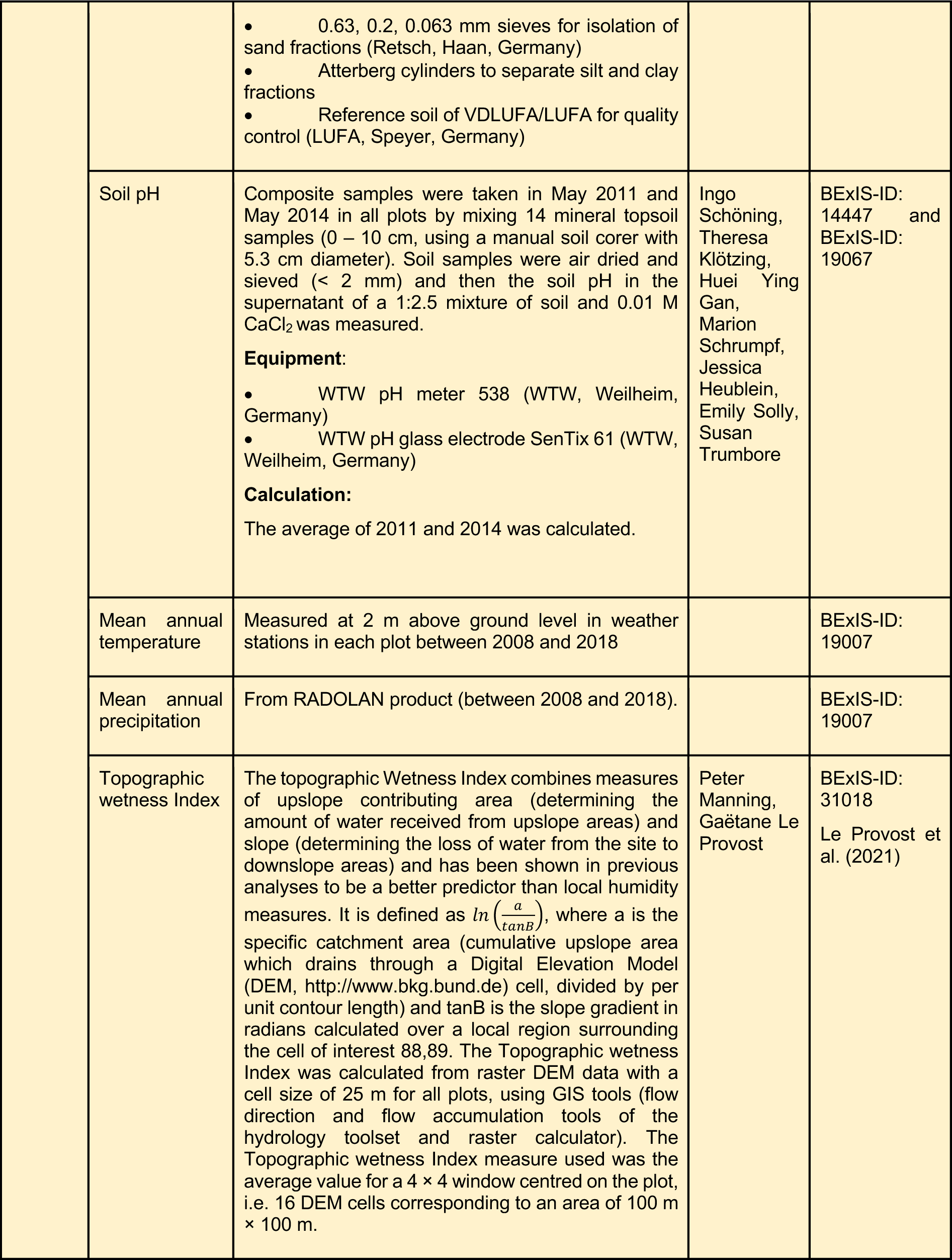
Details to the variables and covariates: Measurement and reference. Each dataset has an ID within the Biodiversity Exploratory project. The dataset can be found using the ID in the Biodiversity Exploratories Information System (BExIS). It is listed under reference. Most biodiversity data were synthesized in BExIS dataset 31206.

**Table S2.** Species of biodiversity with conservation value found in the study area

| List of plant forest specialists from Schmidt et al. (2011) present in the study area (n = 113) |  | List of bird forest specialists from Gregory et al. (2007) and Tucker and Evans (1997) present in the study area (n = 33) | List of bird species from Grüneberg et al. (2016) with conservation status critically endangered, endangered, vulnerable present in the study area (n = 6) |
| --- | --- | --- | --- |
| *Abies alba*<br>*Actaea spicata*<br>*Adoxa moschatellina*<br>*Allium ursinum*<br>*Anemone nemorosa*<br>*Anemone ranunculoides*<br>*Anthriscus sylvestris*<br>*Arctium nemorosum*<br>*Arum maculatum*<br>*Aruncus dioicus*<br>*Asarum europaeum*<br>*Athyrium filix-femina*<br>*Brachypodium sylvaticum*<br>*Calamagrostis arundinacea*<br>*Campanula trachelium*<br>*Cardamine bulbifera*<br>*Cardamine flexuosa*<br>*Cardamine impatiens*<br>- *Carex alba* *Carex digitata*<br>*Carex montana*<br>*Carex remota*<br>*Carex sylvatica*<br>*Carpinus betulus*<br>*Cephalanthera damasonium*<br>*Cephalanthera rubra*<br>*Chaerophyllum temulum*<br>*Chrysosplenium alternifolium*<br>*Convallaria majalis*<br>*Dactylis polygama*<br>*Daphne mezereum*<br>*Digitalis purpurea*<br>*Dryopteris dilatata*<br>*Dryopteris filix-mas*<br>*Elymus caninus*<br>*Epilobium angustifolium*<br>*Epilobium montanum*<br>*Euphorbia amygdaloides*<br>*Euphorbia dulcis*<br>*Fagus sylvatica*<br>*Festuca altissima*<br>*Festuca gigantea*<br>*Fragaria vesca*<br>*Gagea lutea*<br>*Galium odoratum*<br>*Galium rotundifolium*<br>*Galium sylvaticum*<br>*Gymnocarpium dryopteris*<br>*Hedera helix*<br>*Helleborus foetidus*<br>*Hepatica nobilis*<br>*Hordelymus europaeus*<br>*Hypericum hirsutum*<br>*Impatiens noli-tangere*<br>*Impatiens parviflora*<br>*Larix decidua*<br>*Lathraea squamaria* | *Lathyrus vernus*<br>*Lilium martagon*<br>*Listera ovata*<br>*Lonicera xylosteum*<br>*Luzula luzuloides*<br>*Luzula pilosa*<br>*Luzula sylvatica*<br>*Lysimachia nemorum*<br>*Maianthemum bifolium*<br>*Melampyrum pratense*<br>*Melica nutans*<br>*Melica uniflora*<br>*Mercurialis perennis*<br>*Milium effusum*<br>*Moehringia trinervia*<br>*Myosotis sylvatica*<br>*Neottia nidus-avis*<br>*Oxalis acetosella*<br>*Paris quadrifolia*<br>*Phyteuma spicatum*<br>*Poa chaixii*<br>*Poa nemoralis*<br>*Polygonatum multiflorum*<br>*Polygonatum verticillatum*<br>*Polypodium vulgare*<br>*Potentilla sterilis*<br>*Prenanthes purpurea*<br>*Pseudotsuga menziesii*<br>*Pteridium aquilinum*<br>*Pulmonaria obscura*<br>*Quercus rubra*<br>*Ranunculus lanuginosus*<br>*Ribes alpinum*<br>*Rubus idaeus*<br>*Rumex sanguineus*<br>*Sambucus racemosa*<br>*Sanicula europaea*<br>*Senecio ovatus*<br>*Senecio sylvaticus*<br>*Solanum dulcamara*<br>*Sorbus aria*<br>*Sorbus torminalis*<br>*Stachys alpina*<br>*Stachys sylvatica*<br>*Stellaria holostea*<br>*Stellaria nemorum*<br>*Taxus baccata*<br>*Tilia cordata*<br>*Tilia platyphyllos*<br>*Torilis japonica*<br>*Ulmus glabra*<br>*Ulmus laevis*<br>*Veronica montana*<br>*Vicia sylvatica*<br>*Vinca minor*<br>*Viscum album* | *Coccothraustes coccothraustes*<br>*Cyanistes caeruleus*<br>*Dendrocopos minor*<br>*Dryocopus martius*<br>*Ficedula albicollis*<br>*Ficedula hypoleuca*<br>*Fringilla montifringilla*<br>*Garrulus glandarius*<br>*Hippolais icterina*<br>*Jynx torquilla*<br>*Lullula arborea*<br>*Luscinia megarhynchos*<br>*Muscicapa striata*<br>*Oriolus oriolus*<br>*Periparus ater*<br>*Phoenicurus phoenicurus*<br>*Phylloscopus collybita*<br>*Phylloscopus sibilatrix*<br>*Picus canus*<br>*Picus viridis*<br>*Poecile montanus*<br>*Poecile palustris*<br>*Prunella modularis*<br>*Pyrrhula pyrrhula*<br>*Regulus regulus*<br>*Sitta europaea*<br>*Sylvia borin*<br>*Certhia familiaris*<br>*Certhia brachydactyla*<br>*Carduelis spinus*<br>*Carduelis flammea*<br>*Bonasa bonasia*<br>*Anthus trivialis* | *Anthus trivialis*<br>*Ficedula hypoleuca*<br>*Jynx torquilla*<br>*Picus viridis*<br>*Pernis apivorus*<br>*Sturnus vulgaris* |

**Table S3.** Standardised effect size +/− confidence interval and p-values for all response variables.

| Explanatory variables | C Increment | Tree stocks | Soil stocks | Plant specialists richness | Bird specialists richness | Red list |
| --- | --- | --- | --- | --- | --- | --- |
| (Intercept) | -5.68e-03±0.09<br>p = 0.952 | **0.15±0.06**<br>**p = 0.016** | 0.0±0.10<br>p = 0.975 | -0.10±0.09<br>p = 0.244 | 0.08±0.09<br>p = 0.401 | **-0.74±0.15**<br>**p < 0.001** |
| Mean DBH | -0.13±0.09<br>p = 0.144 | **0.65±0.06**<br>**p < 0.001** | -0.01±0.09<br>p = 0.912 | 0.06±0.08<br>p = 0.449 | **0.38±0.09**<br>**p < 0.001** | 0.19±0.12<br>p = 0.126 |
| Deadwood input (sqrt) | -0.04±0.08<br>p = 0.600 | **0.13±0.05**<br>**p = 0.009** | -0.04±0.08<br>p = 0.646 | 0.07±0.07<br>p = 0.319 | 0.11±0.08<br>p = 0.144 | -0.05±0.13<br>p = 0.684 |
| Mixture | -0.09±0.09<br>p = 0.287 | -0.03±0.06<br>p = 0.641 | 0.17±0.09 p =<br>0.066 | **-0.16±0.08**<br>**p = 0.055** | **0.18±0.09**<br>**p = 0.038** | 0.0±0.14<br>p = 0.954 |
| Uneven-aged | **-0.37±0.09**<br>**p < 0.001** | **0.26±0.06**<br>**p < 0.001** | -0.10±0.09 p =<br>0.291 | **-0.23±0.08**<br>**p = 0.005** | 0.0±0.08<br>p = 0.978 | **0.29±0.12**<br>**p = 0.014** |
| Genus (Oak) | -0.01±0.34<br>p = 0.974 | -0.23±0.22<br>p = 0.292 | 0.02±0.37<br>p = 0.958 | 0.21±0.33<br>p = 0.517 | 0.08±0.34<br>p = 0.825 | **0.82±0.39**<br>**p = 0.036** |
| Genus (Pine) | -0.12±0.25<br>p = 0.626 | **-0.75±0.16**<br>**p < 0.001** | -0.11±0.27<br>p = 0.676 | -0.26±0.24<br>p = 0.279 | **-0.59±0.25**<br>**p = 0.018** | -0.24±0.46<br>p = 0.603 |
| Genus (Spruce) | 0.19±0.26<br>p = 0.461 | **-0.35±0.17**<br>**p = 0.041** | 0.09±0.28<br>p = 0.756 | **1.10±0.25**<br>**p < 0.001** | -0.07±0.26<br>p = 0.796 | -0.90±0.61<br>p = 0.144 |
| Observations | 150 | 150 | 150 | 150 | 150 | 150 |

| Explanatory variables | Bacteria richness | Fungi richness | Protist richness | OOmycota richness | Nematode richness | Mollusca richness | Plant richness | Arthropod richness | Bird richness |
| --- | --- | --- | --- | --- | --- | --- | --- | --- | --- |
| (Intercept) | **0.25±0.09**<br>**p = 0.007** | 0.0±0.10<br>p = 0.941 | 0.18±0.09<br>p = 0.058 | 0.05±0.10<br>p = 0.604 | 0.19±0.10 p =<br>0.054 | 0.19±0.10<br>p = 0.058 | **-0.25±0.08**<br>**p = 0.003** | **1.33±0.03**<br>**p < 0.001** | 0.0±0.10<br>p = 0.928 |
| Mean DBH | 0.12±0.08<br>p = 0.146 | **0.18±0.09**<br>**p = 0.053** | **0.21±0.09**<br>**p = 0.016** | **0.19±0.09**<br>**p = 0.041** | 0.14±0.09 p =<br>0.119 | 0.0±0.09 p =<br>0.971 | -0.06±0.08<br>p = 0.423 | 0.04±0.03<br>p = 0.178 | **0.28±0.09**<br>**p = 0.002** |
| Deadwood volume sqrt | 0.14±0.08<br>p = 0.068 | -0.10±0.08<br>p = 0.243 | 0.14±0.08<br>p = 0.074 | 0.02±0.08<br>p = 0.839 | 0.08±0.08 p =<br>0.318 | 0.14±0.08<br>p = 0.099 | 0.04±0.07<br>p = 0.587 | 0.0±0.03<br>p = 0.750 | 0.10±0.08 p =<br>0.231 |
| Mixture continuous | -0.02±0.08<br>p = 0.768 | **0.29±0.09**<br>**p = 0.002** | 0.06±0.09<br>p = 0.482 | 0.11±0.09<br>p = 0.231 | 0.08±0.09 p =<br>0.393 | 0.11±0.09<br>p = 0.255 | -0.12±0.08<br>p = 0.118 | -0.01±0.03<br>p = 0.653 | 0.15±0.09 p =<br>0.094 |
| unevenaged | 0.0±0.08<br>p = 0.940 | -0.04±0.09<br>p = 0.674 | -0.04±0.08<br>p = 0.664 | -0.06±0.09<br>p = 0.508 | **-0.18±0.09**<br>**p = 0.045** | -0.08±0.09<br>p = 0.372 | **-0.19±0.08**<br>**p = 0.015** | -0.03±0.03<br>p = 0.243 | 0.02±0.09 p =<br>0.806 |
| Genus (Oak) | 0.07±0.33<br>p = 0.829 | -0.23±0.36<br>p = 0.522 | 0.28±0.34<br>p = 0.401 | -0.16±0.37<br>p = 0.662 | -0.32±0.36<br>p = 0.372 | -0.46±0.39<br>p = 0.240 | **0.62±0.30**<br>**p = 0.043** | **-0.25±0.12**<br>**p = 0.031** | 0.45±0.35 p =<br>0.201 |
| Genus (Pine) | **-0.89±0.24**<br>**p < 0.001** | 0.19±0.26<br>p = 0.465 | **-1.14±0.25**<br>**p < 0.001** | -0.21±0.27<br>p = 0.432 | **-0.69±0.26**<br>**p = 0.009** | **-0.68±0.30**<br>**p = 0.024** | 0.25±0.22<br>p = 0.260 | 0.04±0.08<br>p = 0.674 | -0.40±0.25 p =<br>0.121 |
| Genus (Spruce) | **-1.24±0.25**<br>**p < 0.001** | -0.16±0.27<br>p = 0.565 | -0.49±0.26<br>p = 0.064 | -0.15±0.29<br>p = 0.611 | **-0.73±0.27**<br>**p = 0.008** | **-0.85±0.28**<br>**p = 0.002** | **1.59±0.23**<br>**p < 0.001** | -0.09±0.09<br>p = 0.327 | 0.30±0.27 p =<br>0.274 |
| Observations | 150 | 150 | 147 | 149 | 149 | 139 | 150 | 147 | 149 |

**Figure S1.**
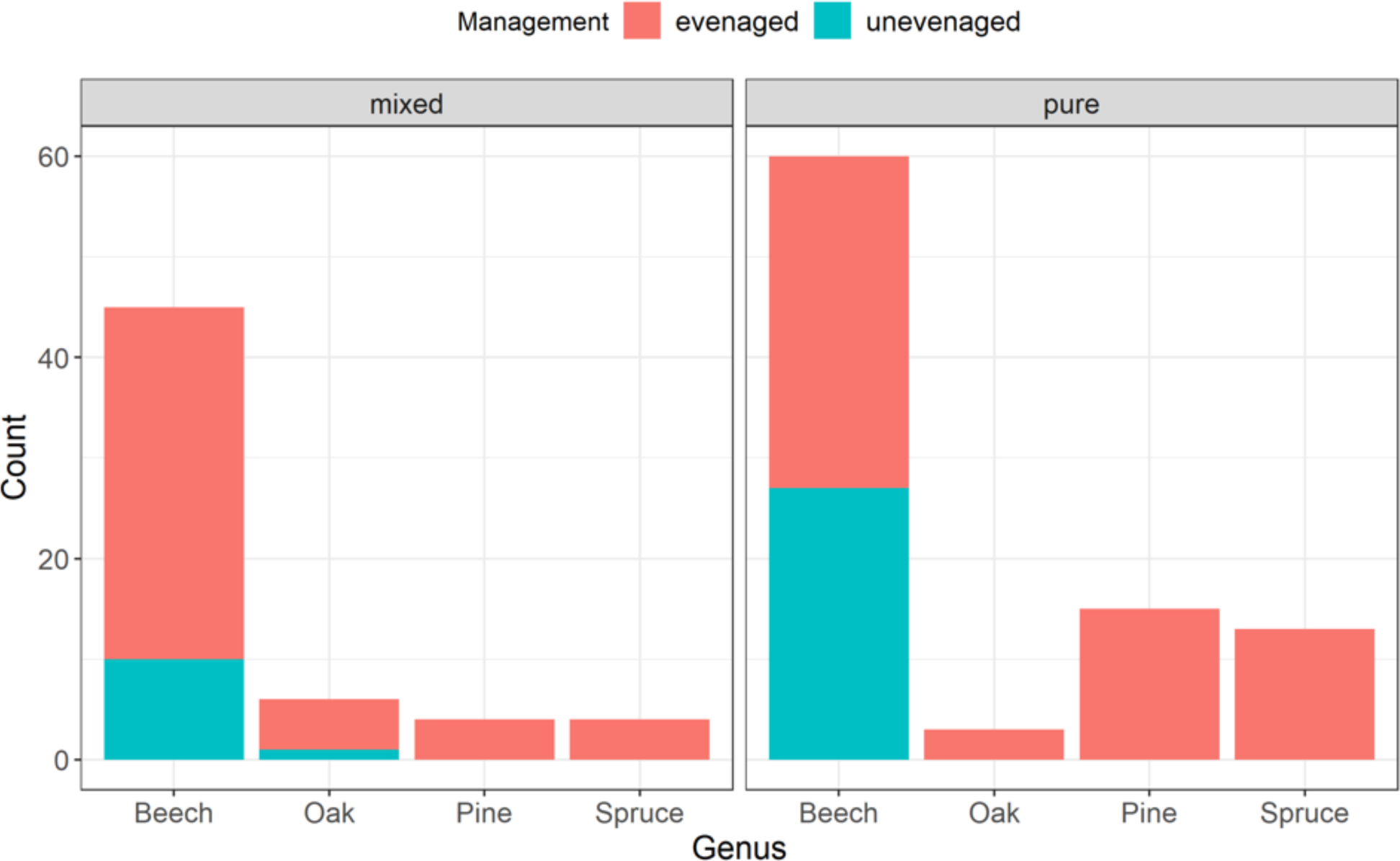
Distribution of management types, dominant species across mixed and pure forests in the study plots. The management type ‘uneven aged’ was only present in forests with the main tree species beech or oak.

**Figure S2.**
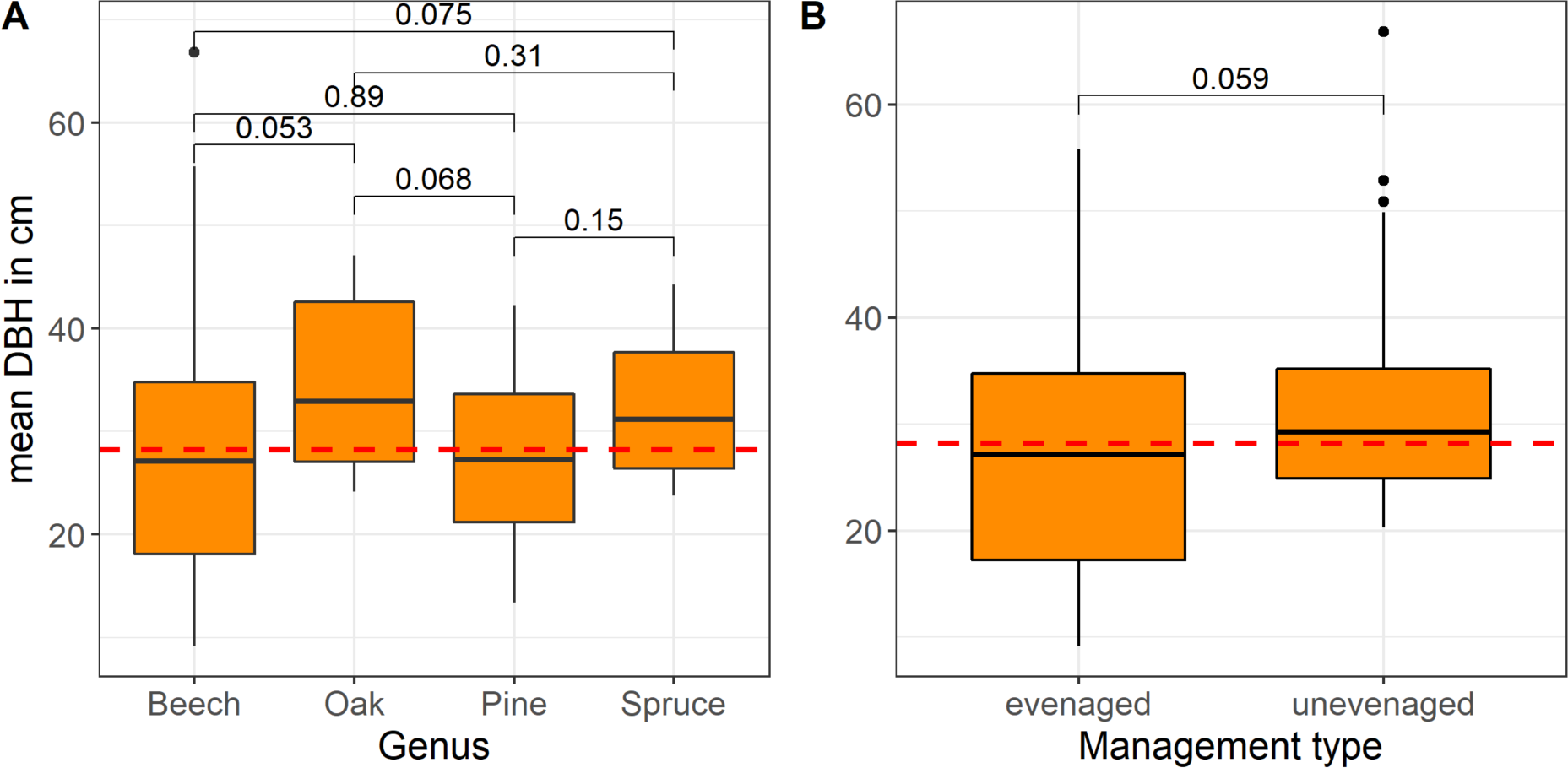
Variation of mean DBH across dominant genera and management type in the study plots; p-values for t-test pairwise comparisons are shown.

**Figure S3.**
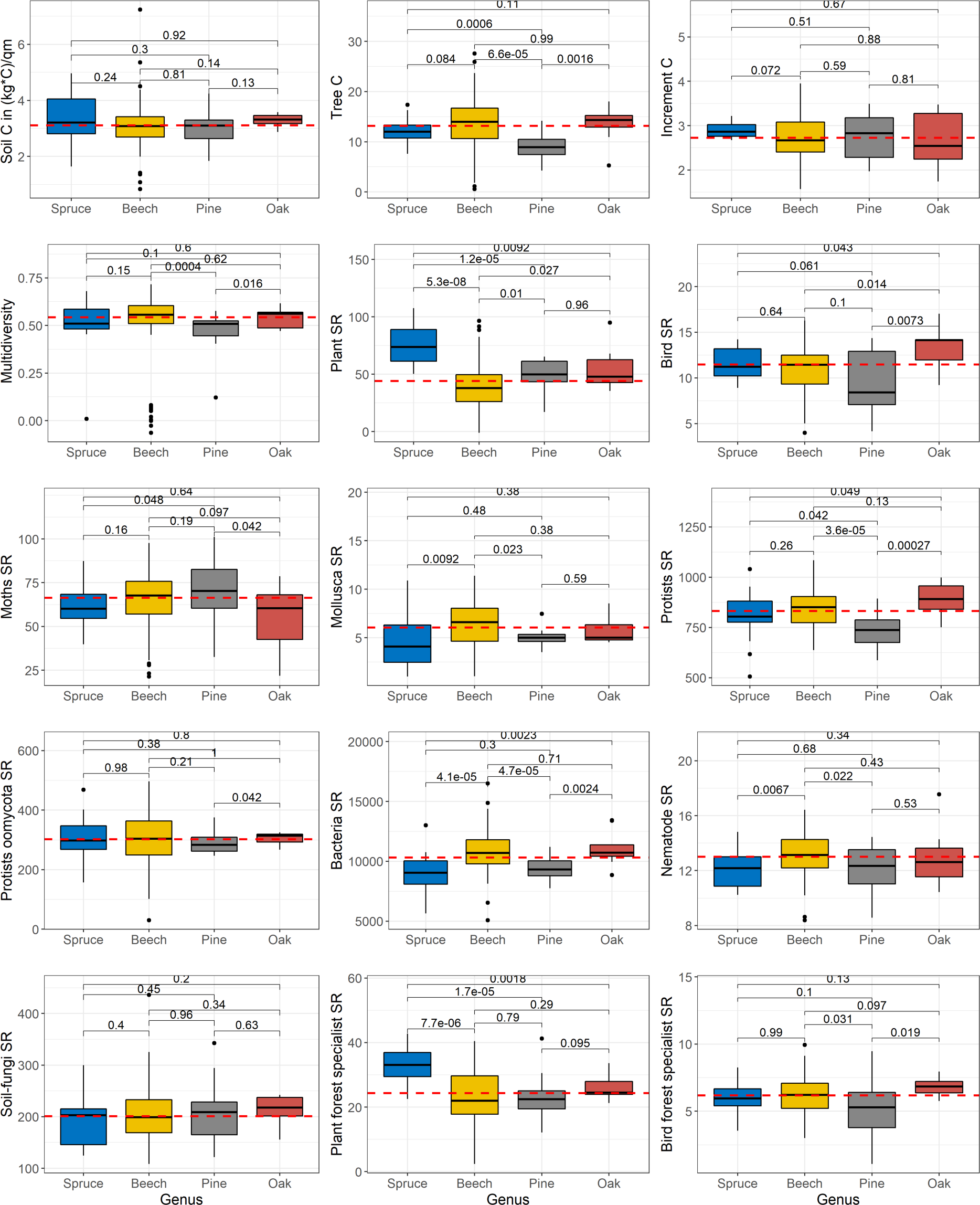
Variation of response variables across dominant species. P values for pairwise t-test comparisons are shown. The red lines mark the individual median of each group. The p-values between each genus are shown.

## Notes

### Competing Interest Statement

The authors have declared no competing interest.

### Summary of Updates

Some minor changes to the abstract and discussion toning down some recommendations Adding some authors' ORCID Adding a few references in the intro

## References

Abele, S. E., Macdonald, S. E., & Spence, J. R. (2014). Cover type, environmental characteristics, and conservation of terrestrial gastropod diversity in boreal mixedwood forests. Canadian Journal of Forest Research, 44(1), 36–44. 10.1139/cjfr-2013-0210

Allan, E., Bossdorf, O., Dormann, C. F., Prati, D., Gossner, M. M., Tscharntke, T., Blüthgen, N., Bellach, M., Birkhofer, K., Boch, S., Böhm, S., Börschig, C., Chatzinotas, A., Christ, S., Daniel, R., Diekötter, T., Fischer, C., Friedl, T., Glaser, K., … Fischer, M. (2014). Interannual variation in land-use intensity enhances grassland multidiversity. Proceedings of the National Academy of Sciences, 111(1), 308–313. 10.1073/pnas.1312213111

Ammer, C. (2019). Buchenverjüngung unter Fichtenschirm – Wachstum und Kohlenstoff-speicherleistung von Umbaubeständen bei langfristigen Verjüngungs-gängen. ALLGEMEINE FORST UND JAGDZEITUNG, 190(3–14), 73–89. 10.23765/afjz0002039

Ammer, C., Bickel, E., & Kölling, C. (2008). Converting Norway spruce stands with beech—A review of arguments and techniques. Austrian Journal of Forest Science, 125, 3–26.

Asbeck, T., Sabatini, F., Augustynczik, A. L. D., Basile, M., Helbach, J., Jonker, M., Knuff, A., & Bauhus, J. (2021). Biodiversity response to forest management intensity, carbon stocks and net primary production in temperate montane forests. Scientific Reports, 11(1), Article 1. 10.1038/s41598-020-80499-4

Birdsey, R. A., Plantinga, A. J., & Heath, L. S. (1993). Past and prospective carbon storage in United States forests. Forest Ecology and Management, 58(1–2), 33–40. 10.1016/0378-1127(93)90129-B

BMEL. (2018). Bundeswaldinventur: Der Wald in Deutschland: Ausgewählte Ergebnisse der dritten Bundeswaldinventur (3; p. 56). Bundesministerium für Ernährung und Landwirtschaft. https://www.bundeswaldinventur.de/dritte-bundeswaldinventur-2012/waldland-deutschland-waldflaeche-konstant/

Boch, S., Prati, D., Müller, J., Socher, S., Baumbach, H., Buscot, F., Gockel, S., Hemp, A., Hessenmöller, D., Kalko, E. K. V., Linsenmair, K. E., Pfeiffer, S., Pommer, U., Schöning, I., Schulze, E.-D., Seilwinder, C., Weisser, W. W., Wells, K., & Fischer, M. (2013). High plant species richness indicates management-related disturbances rather than the conservation status of forests. Basic and Applied Ecology, 14(6), 496–505. 10.1016/j.baae.2013.06.001

Bohn, U., Zazanashvili, N., & Nakhutsrishvili, G. (2007). The Map of the Natural Vegetation of Europe and its application in the Caucasus Ecoregion. 175, 11.

Bonan, G. B. (2008). Forests and Climate Change: Forcings, Feedbacks, and the Climate Benefits of Forests. Science, 320(5882), 1444–1449. 10.1126/science.1155121

Brändle, M., & Brandl, R. (2001). Species richness of insects and mites on trees: Expanding Southwood. Journal of Animal Ecology, 70(3), 491–504. 10.1046/j.1365-2656.2001.00506.x

Brockerhoff, E., Jactel, H., Parrotta, J., Quine, C., & Sayer, J. (2008). Plantation Forests and Biodiversity: Oxymoron or Opportunity? Biodiversity and Conservation, 17, 925–951. 10.1007/s10531-008-9380-x

Brunet, J., Fritz, Ö., & Richnau, G. (2010). Biodiversity in European beech forests – a review with recommendations for sustainable forest management. ECOLOGICAL BULLETINS, 19.

Budde, S., Schmidt, W., & Weckesser, M. (2011). Impact of the admixture of European beech (Fagus sylvatica L.) on plant species diver-sity and naturalness of conifer stands in Lower Saxony. 13.

Carvalho-Santos, C., Sousa-Silva, R., Gonçalves, J., & Honrado, J. P. (2016). Ecosystem services and biodiversity conservation under forestation scenarios: Options to improve management in the Vez watershed, NW Portugal. Regional Environmental Change, 16(6), 1557–1570. 10.1007/s10113-015-0892-0

Charbonnier, Y., Gaüzère, P., van Halder, I., Nezan, J., Barnagaud, J.-Y., Jactel, H., & Barbaro, L. (2016). Deciduous trees increase bat diversity at stand and landscape scales in mosaic pine plantations. Landscape Ecology, 31(2), 291–300. 10.1007/s10980-015-0242-0

Corlett, R. T. (2020). Safeguarding our future by protecting biodiversity. Plant Diversity, 42(4), 221–228. 10.1016/j.pld.2020.04.002

de Lima, R. A. F., Oliveira, A. A., Pitta, G. R., de Gasper, A. L., Vibrans, A. C., Chave, J., ter Steege, H., & Prado, P. I. (2020). The erosion of biodiversity and biomass in the Atlantic Forest biodiversity hotspot. Nature Communications, 11(1), 6347. 10.1038/s41467-020-20217-w

DFWR. (2022). Forest facts—German forestry—300 yrs of sustainability campaign. Forest Facts - German Forestry - 300 Yrs of Sustainability Campaign. https://www.forstwirtschaft-in-deutschland.de/german-forestry/forest-facts/?L=1

Dittrich, S., Jacob, M., Bade, C., Leuschner, C., & Hauck, M. (2014). The significance of deadwood for total bryophyte, lichen, and vascular plant diversity in an old-growth spruce forest. Plant Ecology, 215(10), 1123–1137. 10.1007/s11258-014-0371-6

Dörfler, I., Cadotte, M. W., Weisser, W. W., Müller, J., Gossner, M. M., Heibl, C., Bässler, C., Thorn, S., & Seibold, S. (2020). Restoration-oriented forest management affects community assembly patterns of deadwood-dependent organisms. Journal of Applied Ecology, 57(12), 2429– 2440. 10.1111/1365-2664.13741

Dormann, C. F., Bagnara, M., Boch, S., Hinderling, J., Janeiro-Otero, A., Schäfer, D., Schall, P., & Hartig, F. (2020). Plant species richness increases with light availability, but not variability, in temperate forests understorey. BMC Ecology, 20(1), 43. 10.1186/s12898-020-00311-9

Edelmann, P., Ambarlı, D., Gossner, M. M., Schall, P., Ammer, C., Wende, B., Schulze, E.-D., Weisser, W. W., & Seibold, S. (2022). Forest management affects saproxylic beetles through tree species composition and canopy cover. Forest Ecology and Management, 524, 120532. 10.1016/j.foreco.2022.120532

European Environment Agency. (2016). European forest ecosystems: State and trends. Publications Office. https://data.europa.eu/doi/10.2800/964893

FAO and UNEP. (2020). The State of the World’s Forests 2020. In brief: Forests, biodiversity and people. FAO and UNEP. 10.4060/ca8985en

Felipe-Lucia, M. R., Soliveres, S., Penone, C., Manning, P., van der Plas, F., Boch, S., Prati, D., Ammer, C., Schall, P., Gossner, M. M., Bauhus, J., Buscot, F., Blaser, S., Blüthgen, N., de Frutos, A., Ehbrecht, M., Frank, K., Goldmann, K., Hänsel, F., … Allan, E. (2018). Multiple forest attributes underpin the supply of multiple ecosystem services. Nature Communications, 9(1), 4839. 10.1038/s41467-018-07082-4

Felton, A., Lindbladh, M., Brunet, J., & Fritz, Ö. (2010). Replacing coniferous monocultures with mixed-species production stands: An assessment of the potential benefits for forest biodiversity in northern Europe. Forest Ecology and Management, 260(6), 939–947. 10.1016/j.foreco.2010.06.011

Fiore-Donno, A. M., Richter-Heitmann, T., & Bonkowski, M. (2020). Contrasting Responses of Protistan Plant Parasites and Phagotrophs to Ecosystems, Land Management and Soil Properties. Frontiers in Microbiology, 11. https://www.frontiersin.org/articles/10.3389/fmicb.2020.01823

Fischer, M., Bossdorf, O., Gockel, S., Hänsel, F., Hemp, A., Hessenmöller, D., Korte, G., Nieschulze, J., Pfeiffer, S., Prati, D., Renner, S., Schöning, I., Schumacher, U., Wells, K., Buscot, F., Kalko, E. K. V., Linsenmair, K. E., Schulze, E.-D., & Weisser, W. W. (2010). Implementing large-scale and long-term functional biodiversity research: The Biodiversity Exploratories. Basic and Applied Ecology, 11(6), 473–485. 10.1016/j.baae.2010.07.009

Führer, E. (2000). Forest functions, ecosystem stability and management. Forest Ecology and Management, 10.

Glöer, P. (2017). *Süsswassermollusken: Ein Bestimmungsschlüssel für die Bundesrepublik Deutschland* (15. korrigierte Auflage). Deutscher Jugendbund für Naturbeobachtung (DJN).

Gossner, M. M. (2013). Die Bedeutung von großskaligen Biodiversitätsstudien an Arthropoden am Beispiel der Biodiversitätsexploratorien. 17.

Gregory, R. D., Vorisek, P., Van Strien, A., Gmelig Meyling, A. W., Jiguet, F., Fornasari, L., Reif, J., Chylarecki, P., & Burfield, I. J. (2007). Population trends of widespread woodland birds in Europe: Population trends among widespread woodland birds. Ibis, 149, 78–97. 10.1111/j.1474-919X.2007.00698.x

Grove, S. J. (2002). Saproxylic Insect Ecology and the Sustainable Management of Forests. Annual Review of Ecology and Systematics, 3, 1–23.

Grüneberg, C., Bauer, H.-G., Haupt, H., Hüppop, O., Ryslavy, T., & Südbeck, P. (2016). Rote Liste der Brutvögel Deutschlands. (52; Berichte zum Vogelschutz, pp. 19–67).

Heinrichs, S., Ammer, C., Mund, M., Boch, S., Budde, S., Fischer, M., Müller, J., Schöning, I., Schulze, E.-D., Schmidt, W., Weckesser, M., & Schall, P. (2019). Landscape-Scale Mixtures of Tree Species are More Effective than Stand-Scale Mixtures for Biodiversity of Vascular Plants, Bryophytes and Lichens. Forests, 10(1), 73. 10.3390/f10010073

Hjältén, J., Joelsson, K., Gibb, H., Work, T., Löfroth, T., & Roberge, J.-M. (2017). Biodiversity benefits for saproxylic beetles with uneven-aged silviculture. Forest Ecology and Management, 402, 37–50. 10.1016/j.foreco.2017.06.064

Huston, M. A., & Marland, G. (2003). Carbon management and biodiversity. Journal of Environmental Management, 67(1), 77–86. 10.1016/S0301-4797(02)00190-1

IPBES. (2019). The global assessment report on Biodiversity and Ecosystem services: Summary for policy-makers. 60.

Johnson, S. E., & Abrams, M. D. (2009). Age class, longevity and growth rate relationships: Protracted growth increases in old trees in the eastern United States. Tree Physiology, 29(11), 1317– 1328. 10.1093/treephys/tpp068

Knoke, T., Ammer, C., Stimm, B., & Mosandl, R. (2008). Admixing broadleaved to coniferous tree species: A review on yield, ecological stability and economics. European Journal of Forest Research, 127(2), 89–101. 10.1007/s10342-007-0186-2

Le Provost, G., Thiele, J., Westphal, C., Penone, C., Allan, E., Neyret, M., van der Plas, F., Ayasse, M., Bardgett, R. D., Birkhofer, K., Boch, S., Bonkowski, M., Buscot, F., Feldhaar, H., Gaulton, R., Goldmann, K., Gossner, M. M., Klaus, V. H., Kleinebecker, T., … Manning, P. (2021). Contrasting responses of above- and belowground diversity to multiple components of land-use intensity. Nature Communications, 12(1), 3918. 10.1038/s41467-021-23931-1

Leidinger, J., Blaschke, M., Ehrhardt, M., Fischer, A., Gossner, M. M., Jung, K., Kienlein, S., Kózak, J., Michler, B., Mosandl, R., Seibold, S., Wehner, K., & Weisser, W. W. (2021). Shifting tree species composition affects biodiversity of multiple taxa in Central European forests. Forest Ecology and Management, 498, 119552. 10.1016/j.foreco.2021.119552

Leidinger, J., Weisser, W. W., Kienlein, S., Blaschke, M., Jung, K., Kozak, J., Fischer, A., Mosandl, R., Michler, B., Ehrhardt, M., Zech, A., Saler, D., Graner, M., & Seibold, S. (2020). Formerly managed forest reserves complement integrative management for biodiversity conservation in temperate European forests. Biological Conservation, 242, 108437. 10.1016/j.biocon.2020.108437

Leuschner, C., & Homeier, J. (2022). Global Forest Biodiversity: Current State, Trends, and Threats. In U. Lüttge, F. M. Cánovas, M.-C. Risueño, C. Leuschner, & H. Pretzsch (Eds.), Progress in Botany Vol. 83 (Vol. 83, pp. 125–159). Springer International Publishing. 10.1007/124_2022_58

Lister, B. C., & Garcia, A. (2018). Climate-driven declines in arthropod abundance restructure a rainforest food web. Proceedings of the National Academy of Sciences, 115(44). 10.1073/pnas.1722477115

Löfroth, T., Birkemoe, T., Shorohova, E., Dynesius, M., Fenton, N. J., Drapeau, P., & Tremblay, J. A. (2023). Deadwood Biodiversity. In M. M. Girona, H. Morin, S. Gauthier, & Y. Bergeron (Eds.), Boreal Forests in the Face of Climate Change (Vol. 74, pp. 167–189). Springer International Publishing. 10.1007/978-3-031-15988-6_6

Manning, P., van der Plas, F., Soliveres, S., Allan, E., Maestre, F. T., Mace, G., Whittingham, M. J., & Fischer, M. (2018). Redefining ecosystem multifunctionality. Nature Ecology & Evolution, 2(3), 427–436. 10.1038/s41559-017-0461-7

Mayer, M., Prescott, C. E., Abaker, W. E. A., Augusto, L., Cécillon, L., Ferreira, G. W. D., James, J., Jandl, R., Katzensteiner, K., Laclau, J.-P., Laganière, J., Nouvellon, Y., Paré, D., Stanturf, J. A., Vanguelova, E. I., & Vesterdal, L. (2020). Tamm Review: Influence of forest management activities on soil organic carbon stocks: A knowledge synthesis. Forest Ecology and Management, 466, 118127. 10.1016/j.foreco.2020.118127

McMurdie, P. J., & Holmes, S. (2013). phyloseq: An R Package for Reproducible Interactive Analysis and Graphics of Microbiome Census Data. PLoS ONE, 8(4), e61217. 10.1371/journal.pone.0061217

Meinzer, F. C., Lachenbruch, B., Dawson, T. E., & Meinzer, F. C. (2011). Size- and age-related changes in tree structure and function. Springer.

Michel, A. K., & Winter, S. (2009). Tree microhabitat structures as indicators of biodiversity in Douglas-fir forests of different stand ages and management histories in the Pacific Northwest, U.S.A. Forest Ecology and Management, 257(6), 1453–1464. 10.1016/j.foreco.2008.11.027

Millennium Ecosystem Assessment. (2005). Ecosystems and human well-being: Synthesis; a report of the Millennium Ecosystem Assessment. Island Press.

Moeslund, J. E., Arge, L., Bøcher, P. K., Dalgaard, T., Ejrnæs, R., Odgaard, M. V., & Svenning, J.-C. (2013). Topographically controlled soil moisture drives plant diversity patterns within grasslands. Biodiversity and Conservation, 22(10), 2151–2166. 10.1007/s10531-013-0442-3

Müller, J., Hothorn, T., Yuan, Y., Seibold, S., Mitesser, O., Rothacher, J., Freund, J., Wild, C., Wolz, M., & Menzel, A. (2023). Weather explains the decline and rise of insect biomass over 34 years. Nature. 10.1038/s41586-023-06402-z

Müller, J., Scherer-Lorenzen, M., Ammer, C., Eisenhauer, N., Seidel, D., Schuldt, B., Biedermann, P., Schmitt, T., Künzer, C., Wegmann, M., Cesarz, S., Peters, M., Feldhaar, H., Steffan-Dewenter, I., Claßen, A., Bässler, C., von Oheimb, G., Fichtner, A., Thorn, S., & Weisser, W. (2021). Enhancing the structural diversity between patches for improving multidiversity and multifunctionality in production forests. (p. 11531 KB, 210 pages) [Application/pdf]. Universität Würzburg. 10.25972/OPUS-29084

Neyret, M., Peter, S., Le Provost, G., Boch, S., Boesing, A. L., Bullock, J. M., Hölzel, N., Klaus, V. H., Kleinebecker, T., Krauss, J., Müller, J., Müller, S., Ammer, C., Buscot, F., Ehbrecht, M., Fischer, M., Goldmann, K., Jung, K., Mehring, M., … Manning, P. (2023). Landscape management strategies for multifunctionality and social equity. Nature Sustainability, 6(4), 391–403. 10.1038/s41893-022-01045-w

Oettel, J., Lapin, K., Kindermann, G., Steiner, H., Schweinzer, K.-M., Frank, G., & Essl, F. (2020). Patterns and drivers of deadwood volume and composition in different forest types of the Austrian natural forest reserves. Forest Ecology and Management, 463, 118016. 10.1016/j.foreco.2020.118016

Oksanen, J., Blanchet, F. G., Friendly, M., Kindt, R., Legendre, P., McGlinn, D., Minchin, P. R., O’Hara, R. B., Simpson, G. L., Solymos, P., Stevens, M. H. H., Szoecs, E., & Wagner, H. (2020). vegan: Community Ecology Package (2.5-7) [Computer software]. https://CRAN.R-project.org/package=vegan

Paillet, Y., Bergès, L., Hjältén, J., Ódor, P., Avon, C., Bernhardt-Römermann, M., Bijlsma, R.-J., De Bruyn, L., Fuhr, M., Grandin, U., Kanka, R., Lundin, L., Luque, S., Magura, T., Matesanz, S., Mészáros, I., Sebastià, M.-T., Schmidt, W., Standovár, T., … Virtanen, R. (2010). Biodiversity Differences between Managed and Unmanaged Forests: Meta-Analysis of Species Richness in Europe. Conservation Biology, 24(1), 101–112. 10.1111/j.1523-1739.2009.01399.x

Paillet, Y., Debaive, N., Archaux, F., Cateau, E., Gilg, O., & Guilbert, E. (2019). Nothing else matters? Tree diameter and living status have more effects than biogeoclimatic context on microhabitat number and occurrence: An analysis in French forest reserves. PLOS ONE, 14(5), e0216500. 10.1371/journal.pone.0216500

Penone, C., Allan, E., Soliveres, S., Felipe-Lucia, M. R., Gossner, M. M., Seibold, S., Simons, N. K., Schall, P., Plas, F., Manning, P., Manzanedo, R. D., Boch, S., Prati, D., Ammer, C., Bauhus, J., Buscot, F., Ehbrecht, M., Goldmann, K., Jung, K., … Fischer, M. (2019). Specialisation and diversity of multiple trophic groups are promoted by different forest features. Ecology Letters, 22(1), 170–180. 10.1111/ele.13182

Pettorelli, N., Graham, N. A. J., Seddon, N., Maria da Cunha Bustamante, M., Lowton, M. J., Sutherland, W. J., Koldewey, H. J., Prentice, H. C., & Barlow, J. (2021). Time to integrate global climate change and biodiversity science-policy agendas. Journal of Applied Ecology, 58(11), 2384– 2393. 10.1111/1365-2664.13985

Pörtner, H.-O., Scholes, R. J., Agard, J., Archer, E., Arneth, A., Bai, X., Barnes, D., Burrows, M., Chan, L., Cheung, W. L. (William), Diamond, S., Donatti, C., Duarte, C., Eisenhauer, N., Foden, W., Gasalla, M. A., Handa, C., Hickler, T., Hoegh-Guldberg, O., … Ngo, H. (2021). Scientific outcome of the IPBES-IPCC co-sponsored workshop on biodiversity and climate change. Zenodo. 10.5281/zenodo.5101125

R Core Team. (2023). *R: A Language and Environment for Statistical Computing* [Computer software]. R Foundation for Statistical Computing. https://www.R-project.org/

Remm, J., Lõhmus, A., & Remm, K. (2006). Tree cavities in riverine forests: What determines their occurrence and use by hole-nesting passerines? Forest Ecology and Management, 221(1–3), 267–277. 10.1016/j.foreco.2005.10.015

Richter, A., Ewald, M., Hemmerling, C., Schöning, I., Bauhus, J., Schall, P., & Ruess, L. (2023). Effects of management intensity, soil properties and region on the nematode communities in temperate forests in Germany. Forest Ecology and Management, 529, 120675. 10.1016/j.foreco.2022.120675

Richter, D. D., Markewitz, D., Trumbore, S. E., & Wells, C. G. (1999). Rapid accumulation and turnover of soil carbon in a re-establishing forest. Nature, 400(6739), 56–58. 10.1038/21867

Ruess, L. (1995). Studies On the Nematode Fauna of an Acid Forest Soil: Spatial Distribution and Extraction. Nematologica, 41(1–4), 229–239. 10.1163/003925995X00198

Russ, J. M., & Montgomery, W. I. (2002). Habitat associations of bats in Northern Ireland: Implications for conservation. Biological Conservation, 108(1), 49–58. 10.1016/S0006-3207(02)00089-7

Sabatini, F. M., de Andrade, R. B., Paillet, Y., Ódor, P., Bouget, C., Campagnaro, T., Gosselin, F., Janssen, P., Mattioli, W., Nascimbene, J., Sitzia, T., Kuemmerle, T., & Burrascano, S. (2019). Trade-offs between carbon stocks and biodiversity in European temperate forests. Global Change Biology, 25(2), 536–548. 10.1111/gcb.14503

Sandström, J., Bernes, C., Junninen, K., Lõhmus, A., Macdonald, E., Müller, J., & Jonsson, B. G. (2019). Impacts of dead wood manipulation on the biodiversity of temperate and boreal forests. A systematic review. Journal of Applied Ecology, 56(7), 1770–1781. 10.1111/1365-2664.13395

Schall, P., Gossner, M. M., Heinrichs, S., Fischer, M., Boch, S., Prati, D., Jung, K., Baumgartner, V., Blaser, S., Böhm, S., Buscot, F., Daniel, R., Goldmann, K., Kaiser, K., Kahl, T., Lange, M., Müller, J., Overmann, J., Renner, S. C., … Ammer, C. (2018). The impact of even-aged and uneven-aged forest management on regional biodiversity of multiple taxa in European beech forests. Journal of Applied Ecology, 55(1), 267–278. 10.1111/1365-2664.12950

Schall, P., Heinrichs, S., Ammer, C., Ayasse, M., Boch, S., Buscot, F., Fischer, M., Goldmann, K., Overmann, J., Schulze, E., Sikorski, J., Weisser, W. W., Wubet, T., & Gossner, M. M. (2020). Can multi-taxa diversity in European beech forest landscapes be increased by combining different management systems? Journal of Applied Ecology, 57(7), 1363–1375. 10.1111/1365-2664.13635

Schall, P., Schulze, E.-D., Fischer, M., Ayasse, M., & Ammer, C. (2018). Relations between forest management, stand structure and productivity across different types of Central European forests. Basic and Applied Ecology, 32, 39–52. 10.1016/j.baae.2018.02.007

Schmidt, M., Kriebitzsch, W.-U., & Ewald, J. (Eds.). (2011). Waldartenlisten der Farn-und Blütenpflanzen, Moose und Flechten Deutschlands. Bundesamt für Naturschutz (BfN).

Schulze, E. D. (2018). Effects of forest management on biodiversity in temperate deciduous forests: An overview based on Central European beech forests. Journal for Nature Conservation, 43, 213–226. 10.1016/j.jnc.2017.08.001

Scott, N. A., & Brown, S. (2008). Measuring the Decomposition of Down Dead-Wood. In C. M. Hoover (Ed.), Field Measurements for Forest Carbon Monitoring (pp. 113–126). Springer Netherlands. 10.1007/978-1-4020-8506-2_9

Seddon, N., Turner, B., Berry, P., Chausson, A., & Girardin, C. A. J. (2019). Grounding nature-based climate solutions in sound biodiversity science. Nature Climate Change, 9(2), 84–87. 10.1038/s41558-019-0405-0

Seibold, S., Bässler, C., Brandl, R., Fahrig, L., Förster, B., Heurich, M., Hothorn, T., Scheipl, F., Thorn, S., & Müller, J. (2017). An experimental test of the habitat-amount hypothesis for saproxylic beetles in a forested region. Ecology, 98(6), 1613–1622.

Seidl, R., Schelhaas, M.-J., Rammer, W., & Verkerk, P. J. (2014). Increasing forest disturbances in Europe and their impact on carbon storage. Nature Climate Change, 4(9), 806–810. 10.1038/nclimate2318

Senf, C., & Seidl, R. (2021). Persistent impacts of the 2018 drought on forest disturbance regimes in Europe. Biogeosciences, 18(18), 5223–5230. 10.5194/bg-18-5223-2021

Siitonen, J. (2001). Forest Management, Coarse Woody Debris and Saproxylic Organisms: Fennoscandian Boreal Forests as an Example. Ecological Bulletins, 49, 11–41.

Sikorski, J., Baumgartner, V., Birkhofer, K., Boeddinghaus, R. S., Bunk, B., Fischer, M., Fösel, B. U., Friedrich, M. W., Göker, M., Hölzel, N., Huang, S., Huber, K. J., Kandeler, E., Klaus, V. H., Kleinebecker, T., Marhan, S., von Mering, C., Oelmann, Y., Prati, D., … Overmann, J. (2022). The Evolution of Ecological Diversity in Acidobacteria. Frontiers in Microbiology, 13. https://www.frontiersin.org/journals/microbiology/articles/10.3389/fmicb.2022.715637

Simons, N. K., Felipe-Lucia, M. R., Schall, P., Ammer, C., Bauhus, J., Blüthgen, N., Boch, S., Buscot, F., Fischer, M., Goldmann, K., Gossner, M. M., Hänsel, F., Jung, K., Manning, P., Nauss, T., Oelmann, Y., Pena, R., Polle, A., Renner, S. C., … Weisser, W. W. (2021). National Forest Inventories capture the multifunctionality of managed forests in Germany. Forest Ecosystems, 8(1), 5. 10.1186/s40663-021-00280-5

Six, J., Callewaert, P., Lenders, S., De Gryze, S., Morris, S. J., Gregorich, E. G., Paul, E. A., & Paustian, K. (2002). Measuring and Understanding Carbon Storage in Afforested Soils by Physical Fractionation. Soil Science Society of America Journal, 66(6), 1981–1987. 10.2136/sssaj2002.1981

Soto-Navarro, C., Ravilious, C., Arnell, A., de Lamo, X., Harfoot, M., Hill, S. L. L., Wearn, O. R., Santoro, M., Bouvet, A., Mermoz, S., Le Toan, T., Xia, J., Liu, S., Yuan, W., Spawn, S. A., Gibbs, H. K., Ferrier, S., Harwood, T., Alkemade, R., … Kapos, V. (2020). Mapping co-benefits for carbon storage and biodiversity to inform conservation policy and action. Philosophical Transactions of the Royal Society B: Biological Sciences, 375(1794), 20190128. 10.1098/rstb.2019.0128

Stephenson, N. L., Das, A. J., Condit, R., Russo, S. E., Baker, P. J., Beckman, N. G., Coomes, D. A., Lines, E. R., Morris, W. K., Rüger, N., Álvarez, E., Blundo, C., Bunyavejchewin, S., Chuyong, G., Davies, S. J., Duque, Á., Ewango, C. N., Flores, O., Franklin, J. F., … Zavala, M. A. (2014). Rate of tree carbon accumulation increases continuously with tree size. Nature, 507(7490), 90–93. 10.1038/nature12914

Stokland, J. N., Siitonen, J., & Jonsson, B. G. (2012). Biodiversity in Dead Wood. Cambridge University Press.

Tucker, G. M., & Evans, M. I. (Eds.). (1997). Habitats for birds in Europe: A conservation strategy for the wider environment. BirdLife International.

Turney, C., Ausseil, A.-G., & Broadhurst, L. (2020). Urgent need for an integrated policy framework for biodiversity loss and climate change. Nature Ecology & Evolution, 4(8), 996–996. 10.1038/s41559-020-1242-2

United Nations. (2021, June 10). Tackling Biodiversity & Climate Crises Together and Their Combined Social Impacts. United Nations Sustainable Development. https://www.un.org/sustainabledevelopment/blog/2021/06/tackling-biodiversity-climate-crises-together-and-their-combined-social-impacts/

van der Plas, F., Manning, P., Soliveres, S., Allan, E., Scherer-Lorenzen, M., Verheyen, K., Wirth, C., Zavala, M. A., Ampoorter, E., Baeten, L., Barbaro, L., Bauhus, J., Benavides, R., Benneter, A., Bonal, D., Bouriaud, O., Bruelheide, H., Bussotti, F., Carnol, M., … Fischer, M. (2016). Biotic homogenization can decrease landscape-scale forest multifunctionality. Proceedings of the National Academy of Sciences, 113(13), 3557–3562. 10.1073/pnas.1517903113

von Lüpke, B., Ammer, C., Bruciamacchie, M., Brunner, A., Ceitel, J., Collet, C., Deuleuze, C., Di Placido, J., Huss, J., Jankovič, J., Kantor, P., Larsen, J. B., Lexer, M., Löf, M., Longauer, R., Madsen, P., Modrzyński, J., Mosandl, R., Pampe, A., … Zientarski, J. (2004). Silvicultural Strategies for Conversion. In H. Spiecker, J. Hansen, E. Klimo, J. P. Skovsgaard, H. Sterba, & K. von Teuffel (Eds.), Norway Spruce Conversion (pp. 121–164). BRILL. 10.1163/9789047412908_009

Vries, W., Reinds, G. J., Posch, M., Sanz-Sanchez, M.-J., Krause, G. H. M., Calatayud, V., Renaud, J.-P., Dupouey, J.-L., Sterba, H., Vel, E. M., Dobbertin, M., Gundersen, P., & Voogd, J. C. H. (2003). Intensive Monitoring of Forest Ecosystems in Europe: Technical Report 2003.

Vuidot, A., Paillet, Y., Archaux, F., & Gosselin, F. (2011). Influence of tree characteristics and forest management on tree microhabitats. Biological Conservation, 144(1), 441–450. 10.1016/j.biocon.2010.09.030

Wagner, S., Fischer, H., & Huth, F. (2011). Canopy effects on vegetation caused by harvesting and regeneration treatments. European Journal of Forest Research, 130(1), 17–40. 10.1007/s10342-010-0378-z

Wehner, K., Renker, C., Simons, N. K., Weisser, W. W., & Blüthgen, N. (2021). Narrow environmental niches predict land-use responses and vulnerability of land snail assemblages. BMC Ecology and Evolution, 21(1), 15. 10.1186/s12862-020-01741-1

Wellbrock, N., Grüneberg, E., Riedel, T., & Polley, H. (2017). Carbon stocks in tree biomass and soils of German forests. Central European Forestry Journal, 63(2–3), 105–112. 10.1515/forj-2017-0013

Welter-Schultes, F. W. (Ed.). (2012). European non-marine molluscs: A guide for species identification = Bestimmungsbuch für europäische Land-und Süsswassermollusken (1. ed). Planet Poster Ed.

Wiese, V. (2016). Die Landschnecken Deutschlands: Finden - erkennen - bestimmen (2., durchgesehene Auflage). Quelle & Meyer Verlag.

Winter, S., & Möller, G. C. (2008). Microhabitats in lowland beech forests as monitoring tool for nature conservation. Forest Ecology and Management, 255(3–4), 1251–1261. 10.1016/j.foreco.2007.10.029

